# How do invasion syndromes evolve? An experimental evolution approach using the ladybird Harmonia axyridis

**DOI:** 10.1101/849968

**Authors:** Julien Foucaud, Ruth A. Hufbauer, Virginie Ravigné, Laure Olazcuaga, Anne Loiseau, Aurélien Ausset, Su Wang, Lian-Sheng Zang, Nicolas Leménager, Ashraf Tayeh, Arthur Weyna, Pauline Gneux, Elise Bonnet, Vincent Dreuilhe, Bastien Poutout, Arnaud Estoup, Benoît Facon

## Abstract

Experiments comparing native to introduced populations or distinct introduced populations to each other show that phenotypic evolution is common and often involves a suit of interacting phenotypic traits. We define such sets of traits that evolve in concert and contribute to the success of invasive populations as an ‘invasion syndrome’. The invasive Harlequin ladybird *Harmonia axyridis* displays such an invasion syndrome with, for instance, females from invasive populations being larger and heavier than individuals from native populations, allocating more resources to reproduction, and spreading reproduction over a longer lifespan. Invasion syndromes could emerge due to selection acting jointly and directly on a multitude of traits, or due to selection on one or a few key traits that drive correlated indirect responses in other traits. Here, we investigated the degree to which the *H. axyridis* invasion syndrome would emerge in response to artificial selection on either female body mass or on age at first reproduction, two traits involved in their invasion syndrome. To further explore the interaction between environmental context and evolutionary change in molding the phenotypic response, we phenotyped the individuals from the selection experiments in two environments, one with abundant food resources and one with limited resources. The two artificial selection experiments show that the number of traits showing a correlated response depends upon the trait undergoing direct selection. Artificial selection on female body mass resulted in few correlated responses and hence poorly reproduced the invasion syndrome. In contrast, artificial selection on age at first reproduction resulted in more widespread phenotypic changes, which nevertheless corresponded only partly to the invasion syndrome. The artificial selection experiments also revealed a large impact of diet on the traits, with effects dependent on the trait considered and the selection regime. Overall, our results indicate that direct selection on multiple traits was likely necessary in the evolution of the *H. axyridis* invasion syndrome. Furthermore, they show the strength of using artificial selection to identify the traits that are correlated in different selective contexts, which represents a crucial first step in understanding the evolution of complex phenotypic patterns, including invasion syndromes.

## Introduction

The field of biological invasions has its own quest for the Holy Grail: establishing a list of traits that can predict invasion success. Traits linked to invasion success and ecological or economic impact are already used in risk assessment to identify potentially harmful species, and then focus efforts to prevent introduction of those species (Kumschick & Richardson 2013). In plants, such risk assessment is commonly based on the evaluation of five characteristics related to reproduction and habitat use (seed mass, chromosome number, native range size, wetland association and maximum height; Schmidt et al. 2012). Unfortunately, these trait-based approaches have met limited success, belying the existence of universal traits that predict invasiveness (Catford et al. 2009, Perkins et al. 2011). Clearly, further refining our understanding of the kinds of traits, and suites of traits, that predict invasion success will move us closer to being able to predict, and prevent, biological invasions. A first step in this process is acknowledging that traits do not exist in isolation from each other; organisms are physiologically integrated units (Ketterson et al. 2009), and organismal traits are constrained by physiological and genetic trade-offs.

Research focused on identifying traits that can predict biological invasions has been happening concurrently with a quickly growing body of research that has revealed biological invasions as crucibles for rapid evolution (Reznick & Ghalambor 2001, Lee et al. 2007, Keller & Taylor 2008, Phillips et al. 2010, Estoup et al. 2016). These studies have shown not only that evolutionary changes can be extremely fast, but that rapid evolution increases the success of invasive species (see for instance Blair & Wolfe 2004, Phillips et al. 2006, Colautti & Lau 2015), and thus likely their economic and ecological impacts as well. Thus, it is now obvious that accounting for evolutionary change is critical for developing robust predictive models of biological invasions as well as sound long-term approaches both for preventing future invasions and for managing existing ones (Estoup et al. 2016, Reznick et al. 2019).

These two bodies of work overlap with experiments that compare native to introduced populations or distinct introduced populations to each other, and show that phenotypic evolution is common (Bossdorf et al. 2005, Dlugosch & Parker 2008, Phillips et al. 2010, Colautti & Lau 2015, Chuang & Peterson 2016) and often involves changes in a suite of interacting phenotypic traits (such as growth, reproduction, dispersal and defense against enemies) rather than in a single trait, (Blair & Wolfe 2004, Phillips et al. 2006, Colautti et al. 2010). These suites of phenotypic changes could be crucial in the evolution of invasive populations, because interacting traits may form opportunities for or constraints to phenotypic change that are different from the evolution of a single trait. Here, we define the set of traits that evolve in concert and contribute to the success of invasive populations as an ‘invasion syndrome’. Understanding how such invasion syndromes evolve within invasive populations is critical to assess the extent to which invasions might be repeatable or predictable, as well as the degree to which such suites of traits can be used in risk assessment and prevention of new invasions.

A powerful biological model for studying rapid evolution associated with invasions is provided by the invasive Harlequin ladybird *Harmonia axyridis* (Roy et al. 2016). Native to Asia, *H. axyridis* was intentionally introduced into many countries as a biological control agent of pest insects and is now considered invasive nearly worldwide (Lombaert et al. 2014). Population size in the invasive range can fluctuate dramatically (Brown et al. 2007), likely leading to strong differences in resource availability, and providing an ecologically relevant environmental axis for experimentally evaluating the evolution of traits involved in the invasion. Additionally, invasive *H. axyridis* exhibits a clear invasive syndrome, involving several evolutionary shifts (reviewed in Tayeh et al. 2012 and Roy et al. 2016). *Harmonia axyridis* females from invasive populations reproduce earlier, allocate more resources to reproduction, and spread reproduction over a longer lifespan. They are larger and heavier than individuals from native populations (Tayeh et al. 2015, and see Table 1).

**Table 1:** Phenotypic changes observed in invasive populations of *H. axyridis* that contribute to the invasion syndrome and the corresponding changes displayed by the experimental lines of the selection experiments achieved in the present study on female body mass (heavy lines) and age at first reproduction (fast lines) using an *ad libitum* nutritional environment for the phenotyping step. See Facon et al. 2011, Tayeh et al. 2012 and Tayeh et al. 2015 for additional details about phenotypic characteristics of invasive *H. axyridis* populations.

| Trait | Natural invasive | Experimental selection |  |
| --- | --- | --- | --- |
|  | Populations | Heavy lines | Fast lines |
| Hatching rate | No change | No change | No change |
| Larval development | Faster | No change | Faster |
| Larval survival | No change | No change | No change |
| Female body mass | Heavier | Heavier | Heavier |
| Age at first reproduction | Earlier | No change | Earlier |
| Fecundity | Higher | No change | Lower |
| Survival | Longer | No change | Shorter |

Additionally, invasive males mate more often and outperform native males in sperm competition (Laugier et al. 2013). Larvae show a higher propensity for cannibalism in invasive populations than in native ones (Tayeh et al. 2014). Finally, invasive populations experience almost none of the inbreeding depression for lifetime performance and generation time suffered by native populations, likely because deleterious alleles were purged during the introduction and invasion process (Facon et al. 2011). Importantly, this suite of evolutionary shifts in morphological, behavioural and life-history traits has been consistently found in different invasive populations originating from distinct source populations, strongly suggesting that their evolution has played a role in the invasion process itself. These evolutionary changes constitute the invasion syndrome of *H. axyridis*. Three potential explanations for this invasion syndrome (see Tayeh et al. 2015 for details) are as follows. First, invasive individuals by chance might have a higher rate of resource acquisition than the native ones (Reznick et al. 2000), providing them with the resources needed to support the shifts in traits that have been observed. Second, possible trade-offs with other (unmeasured) traits could be large in the native area but smaller in the invasive area allowing invasive populations to reallocate resources to growth and reproduction. Third, deleterious mutations could have been purged from invasive *H. axyridis* populations during the course of the invasion. This could enable invasive individuals to exhibit higher fitness than native ones (Facon et al. 2011).

While the existence of an invasion syndrome, such as the one described above for *H. axyridis*, can be well-established, identifying the evolutionary forces that produced it is challenging. The evolution of a given invasion syndrome could be due to a simple selective process acting on a single trait that drives the evolution of other genetically-correlated traits (i.e. correlated responses to selection; e.g. Irwin & Carter 2014). In this case, the syndrome may stem from historical selection (i.e. past selective pressures that have favored integrated combinations of particular traits) or from pleiotropic effects (one or few genes that influence multiple phenotypic traits). If the syndrome relies on historical selection, the genetic correlations could be more easily broken and subsequent evolutionary changes would be relatively unconstrained (Roff 1997) compared to the case where the syndrome relies on pleiotropic effects (Schluter 1996). The evolution of an invasion syndrome could also be linked to complex selection pressures acting jointly and directly on multiple traits (Anderson et al. 2010). Indeed, selection on multiple traits could happen either during the introduction process or during the range expansion within the new range. First, the introduction events act as selective filters that can pick up individuals displaying particular suites of behaviors such as having higher foraging activity or being bolder and more aggressive (Blackburn & Duncan 2001, Chapple et al. 2012). Then, selective pressures experienced during expansion across the invaded region can drive the evolution of distinctive phenotypic traits. An excellent example of this occurs when individuals on an invasion front are assorted by dispersal ability and also experience a lower-density environment than individuals behind the front. This spatial dynamic leads to the evolution of increased dispersal and reproductive rates, and can select for decreases in resources allocated to defense against specialized enemies (Phillips et al. 2010, Chuang & Peterson 2016).

Which of these processes underlies the evolution of invasion syndromes is difficult to infer from natural populations. This limits our ability to pinpoint the evolutionary forces and the genetic bases of the traits involved in the invasion syndromes (Keller & Taylor 2008). Approaches based on experimental evolution and selection provide a means of overcoming limitations inherent to studying wild populations as the evolutionary processes imposed are evident, and the underlying genetic basis of traits can be known (Kawecki et al. 2012, Bataillon et al. 2013). An outstanding example of the use of selection experiments to understand invasions is the study of the transition from saline to freshwater by the marine copepod Eurytemora affinis (Lee et al. 2011, Lee 2016). In replicate laboratory selection experiments, Lee et al. (2011) found that adaptation to freshwater was repeatedly accompanied by evolution of ion-motive enzyme activity and development rate (Lee et al. 2007, Lee 2016). Habitat transition in both natural and laboratory settings in this copepod predictively led to the evolution of a suite of interacting phenotypic traits contributing to invasion success (i.e. an ‘invasion syndrome’).

Importantly, invaders are likely to experience a variety of environments during invasions (including benign and stressful ones), but whether and how environment quality influences the expression of invasion syndromes remains an open question. More specifically, invasive species often initially experience abundant resources (Davis & Pelsor 2001, Tyler et al. 2007, Blumenthal et al. 2009) and escape from predation (Garvey et al. 1994, Roy et al. 2011). As population densities increase, intense intraspecific competition may lead to resource stress (Gioria & Osborne 2014). Resource availability may also change from generation to generation, particularly as densities become high in the introduced range, and thus diet may change with the shifting resource base (Tillberg et al. 2007). It has been shown that diet variation can have huge impacts on trait variances and covariances. For instance, diet reduction may generate trade-offs by prioritizing some physiological functions over others (Royauté et al. 2019).

The main aim of the present study is to understand the evolution of invasion syndromes, and how consistently they are expressed following different selection regimes and in different environmental contexts. More specifically, our goal was to better understand the invasion syndrome of *H. axyridis*. To achieve this goal, we performed two distinct artificial selection experiments, each focused on a phenotypic trait involved in the *H. axyridis* invasion syndrome and we examined the potential joint evolution on other life-history traits. The two focal traits, body mass and age at first reproduction of females, show marked differences between native and invasive populations of *H. axyridis* (see Supplementary Figure 1). Both selection experiments started from individuals sampled in the native range of the species (China). After multiple generations of experimental selection, we measured the selected traits as well as other life-history traits, several of which are also part of the *H. axyridis* invasion syndrome (hatching rate, survival and fecundity). To explore the interaction between the environmental context and the evolutionary changes in molding the phenotypic response, we measured these traits in two environments: one with abundant food resources and one with limited resources (hereafter referred to as *ad libitum* and stress, respectively). Our study addressed the following specific questions: 1) Do body size and age at first reproduction respond to selection? 2) How does selection on these individual traits affect other traits in the invasion syndrome, specifically what are the directions and magnitudes of correlated responses to selection? 3) How are the relationships between phenotypic traits affected by selection on a single trait? And 4) to what extent does the environmental context, here food availability, modify the expression of the phenotypic syndrome that evolved through artificial selection?

## Methods

### Sampling and laboratory rearing conditions

Adult *H. axyridis* individuals were sampled in the native area of the Jilin province, China (43°58’ N, 125°45’ E) in October 2013 and 2015. This population was previously sampled and found to be a source of the first invasive population of *H. axyridis* established in Western North America (N-China3 population from Lombaert et al, 2011). The 2013 sampling was used for artificial selection on female body mass and the 2015 sampling was used for selection on age at first reproduction of females (the first time when females laid eggs). In 2013, collected individuals were first maintained in diapause at 4°C during the whole winter, before gradually breaking diapause in April 2014. Approximately 1,500 individuals were collected, of which approximately 900 survived the winter and were allowed to mate and lay eggs, constituting what we hereafter call base population 1. The individuals collected in October 2015 were not maintained in diapause for a long period, as they were immediately put in conditions to gradually break diapause in November 2015. Approximately 1,500 individuals were collected, of which approximately 1,000 survived, constituting what we hereafter call base population 2. As both populations experienced diapause and showed similar rates of mortality during their diapause, we assumed that different durations of diapause did not induce large differences between the 2013 and 2015 populations. All *H. axyridis* rearing took place at 25°C on 14L:10D light cycle. Unless otherwise stated, *H. axyridis* were fed *ad libitum* with *Ephestia kuhnellia* eggs.

### Heritability estimates of female body mass and age at first reproduction

A full-sib design was used to estimate simultaneously the broad-sense heritabilities of female body mass and age at first reproduction using base population 1. Forty third-generation couples were formed randomly and their eggs were allowed to develop. Of the 30 couples that had offspring, 24 produced 10 or more female offspring that survived to adulthood. These females were used to estimate heritability (total = 485 female offspring from 24 families). The remaining six families were not included because they had too few offspring, which would unnecessarily inflate the variance of our heritability estimations. Every adult female offspring was weighed two days after emergence. Each female was then paired with a male of the same age randomly chosen from base population 1. *Harmonia axyridis* is a multi-copulating species (i.e. males and females can mate multiple times with the same or different individuals during their lives; Laugier et al. 2013). Each female was presented a new male from base population 1 every day for 21 days, in order to limit the effect on reproduction that might be observed if only a single particularly poor or high quality male was used. Age at first reproduction of females was recorded as the number of days between hatching of the female as an adult and the day she first laid a clutch of eggs.

For each trait, broad-sense heritability was computed using intra-class (family) correlation and standard errors were estimated taking into account unequal family sizes using eq. 2.29 from Roff (1997). Because multiple paternity is likely in our experimental set-up, offspring could be full- or half-siblings. This design may thus systematically underestimate true broad-sense heritability, making our estimates of broad-sense heritability estimate conservative. The goal of these estimates of broad-sense heritability was to ensure that the phenotypic traits possessed sufficient additive genetic variance to respond to artificial selection. In addition, we also estimated narrow-sense heritabilities of the selected traits using the mean response to selection per generation, the intensity of selection (in both selection experiments, truncation is at *p* = 0.25 so *i* = 1.27) and the phenotypic variance available at the beginning of the selection experiments (Roff 1997).

### Divergent selection on female body mass

At the eighth generation of base population 1 (referred to as G0 hereafter), the selection experiment on female body mass was initiated. Selection on body mass was carried out in females only, because only female *H. axyridis* evolved larger size during the course of worldwide invasion. Two days after emergence, 700 females were weighed, and 14 experimental lines were established as follows (Supplementary Figure 2A). Four control lines were founded by randomly sampling 50 females across the entire body mass distribution for each line. Five light lines were founded by randomly sampling 50 females from the 50% lower part of the female body mass distribution for each line. Five heavy lines were founded by randomly sampling 50 females from the 50% higher part of the body mass distribution for each line. Each of the 14 experimental lines was supplemented with 50 males randomly chosen from base population 1. Once founded, all lines were maintained independently for the rest of the experimental selection (there was no gene flow between lines; Supplementary Figure 2B). For each subsequent generation, 200 females were weighed two days after emergence for each line and 50 females were chosen to initiate the next generation. For the control lines, 50 random females were chosen, for the light lines, 50 females were chosen from the lightest quartile, and for the heavy lines, 50 females were chosen from the heaviest quartile. For each line, 10 days after emergence, 50 males were randomly chosen from the relevant line and added to the 50 females. Fourteen days after emergence, >70 clutches per line were collected to establish the next generation. While mating and egg-laying were carried out in a single arena per line containing all individuals, we are confident that most individuals of both sexes contributed to next generation because (i) number of collected clutches was generally very high (>200 clutches), (ii) repeated mating was confirmed visually to occur all four days. Collected clutches were distributed into several dozens of small boxes per line (diameter = 55mm; 2 clutches per box), to ensure a balanced contribution of all clutches to the next generation. After hatching, a minimum of 33 boxes per line, each containing 12 larvae from a single hatching box (i.e., minimum n = 396 individuals per line), was reared and fed by adding *E. kuhnellia* eggs *ad libitum* twice a week until emergence of adult individuals. The use of small boxes with a low density of larvae enabled us to limit cannibalism, which otherwise is common in this species (Tayeh et al. 2014) and to prevent an unbalanced contribution of clutches to the next generation. This selection scheme was continued for eight generations.

### Directional selection on the age at first reproduction of females

At the second generation of base population 2, the distribution of the age at first reproduction was estimated for females using 128 couples isolated at emergence. Males were rotated between females each day to limit the effect of males on the timing of oviposition. From this distribution, we determined the numbers of days at which 25% and 75% of females had reproduced and laid their first clutch, respectively. Those numbers (25% quartile = 7 days, 75% quartile = 21 days) were used to determine the day on which eggs were to be collected to establish both selected and control experimental lines. At the third generation, ten experimental lines were established from 240 randomly chosen egg clutches as follows (Supplementary Figure 2C). Seven fast lines were founded by collecting egg clutches that were laid on days 5-7 (the 2 days before the 25% quartile of the distribution of age at first reproduction). Three control lines were founded by collecting eggs laid on days 19-21 (the two days before the 75% quartile). Note that these lines included eggs from the earliest reproducing females as well as eggs from later reproducing females and that all eggs laid prior to day 19 were removed every two days (and hence did not participate in the next generation). Those quartiles were chosen to: (i) ensure sufficient production of females and males to sustain the selection experiment in the fast lines, and (ii) provide a maximum number of individuals with the opportunity to participate in reproduction, while reasonably limiting generation time in the control lines. At each generation, selection proceeded as follows (Supplementary Figure 2D). Each of the egg clutches collected was maintained in a separate 5cm-diameter box and larvae were maintained at a maximal density of 12 individuals per box during their development. Upon emergence, adults of each line were moved into a single cage, and date of the peak of emergence of each experimental line was recorded. For fast lines, new egg clutches were counted and collected each day until 20 clutches were collected. For control lines, new egg clutches were counted but only collected during the 48-hrs period before the 75% quartiles of the distribution of ages (i.e., between days 19-21 after the peak of emergence), except at generation G2 for technical reasons. Each fast and control line was maintained at approximately 100 females and 100 males able to mate freely for the entire selection process. This selection scheme was continued for nine generations.

### Phenotyping of experimental lines

At the end of the selection experiments, we phenotyped individuals of each experimental line using the same protocol for the following traits: hatching rate, larval survival rate, development time, female and male body mass, age at first reproduction of females, female fecundity and adult survival (Supplementary Figure 3). These traits span the life history of *H. axyridis* and five of them are involved in the *H. axyridis* invasion syndrome. Apart from hatching rate, all traits were measured on individuals reared in two environments: either with *ad libitum* food resources or with limited food resources (hereafter *ad libitum* and stressful conditions, respectively). Individuals from the female body mass experiment were phenotyped at G9 and those from the age at first reproduction experiment were phenotyped at G10.

Specifically, females from the generation just before phenotyping (G8 for body mass and G9 for age at first reproduction) were allowed to mate and lay eggs for 24h, 19 days after emergence as adults. For each line, 30 clutches were randomly chosen and their number of eggs were counted. The clutches were then placed in individual 8cm-diameter boxes. Hatching rate was recorded for each clutch three days after collection. For each line, we monitored larval development of groups of 10 larvae under an *ad libitum* food diet (1.5g of *E. kuhnellia* eggs per group, twice a week) and groups of 10 larvae under a stressful food diet (0.5g of *E. kuhnellia* eggs per group, twice a week), maintained in 8cm-diameter boxes. For each of the 14 lines selected for female body mass, we monitored eight groups of 10 larvae fed *ad libitum* and 10 groups of 10 larvae under food stress (for a total of 2,520 larvae). For each of the 10 lines selected for age at first reproduction, we monitored 15 groups of 10 larvae fed *ad libitum* and 15 under food stress (for a total of 3,000 larvae). We recorded the number of surviving larvae, pupae and adults daily for the entire duration of development to adulthood. These data were used to compute development time from egg to adult and larval survival rate. Upon emergence, individuals were sexed and moved to individual boxes. All individuals that survived to adulthood (both males and females) were weighed two days after their emergence. When possible, couples were formed immediately after weighting (or the next day at the latest), by keeping one male and one female in a 5cm-diatemeter box together with the same diet they received as larvae (*ad libitum* diet: 0.35g of *E. kuhnellia* eggs per couple twice a week; stressful diet: 0.1g per couple twice a week). Twenty-four couples were formed per female body mass line per diet and 48 couples were formed per line selected for age at first reproduction (total of 672 and 960 monitored couples, respectively). Females were checked daily for eggs and males were rotated between females to limit the effect of individual males on female fecundity. When eggs were laid, age at first reproduction was recorded and both the female and its current male partner were placed in a large box (10 × 5 × 5cm) containing reproducing couples of the same line and diet, with a maximum of 48 individuals per box. Individuals from each line and diet were gathered in one box and two boxes for the female body mass and age at first reproduction selection experiments, respectively (total = 28 and 40 boxes, respectively). Adults that had already reproduced were fed twice a week, again following a diet that matched their larval and adult stages (*ad libitum* diet: 0.17g of *E. kuhnellia* eggs per individual twice a week; stressful diet: 0.05g per individual twice a week). Once a week, fecundity was measured by counting the number of clutches laid during the last 24h. Twice a week, adult survival was measured by counting and sexing live and dead adults. Fecundity and survival were monitored until the death of all individuals (1,344 and 1,920 individuals followed for body mass and age at first reproduction schemes, respectively).

### Statistical treatments

All data and R code are available at: https://data.inra.fr/dataset.xhtml?persistentId=doi:10.15454/V9XCA2

For each variable (i.e. trait), we performed a Generalized Linear Mixed Model (GLMM) analysis with Laplace approximation (Bolker et al. 2009), using the *lme4* package (Bates et al. 2014) in the R statistical framework (R Core Team). Depending on variable distribution, we ran an LMM for Gaussian variables (i.e., female and male body mass), a binomial GLMM with a logit link for proportion variables (i.e., hatching rate, larval survival rate) and a Poisson GLMM with a log link for Poisson distributed variables (i.e., development time, age at first reproduction, number of eggs laid and age at death of females and males). For all individual variables, we investigated the significance of the following explanatory factors: selection type (selected or control during the experimental selection assay), diet type (*ad libitum* or stressful diet during the phenotyping experiment) and line (the identity of replicate experimental lines during selection). Both selection and diet types were treated as fixed effects, while line was treated as a random effect. To account for over-dispersion in some variables, we added an observation-level random effect to the models when necessary, as recommended by Bolker et al. (2009). We followed a step-by-step procedure of model simplification from the full model (starting with random effects and then fixed effects) based on the Akaike Information Criterion (AIC), and tested the significance of the remaining effects via Likelikood Ratio Tests (LRT). Normality of residuals was checked using quantile-quantile plots. Adjusted means and confidence intervals for each significant fixed effect were calculated using a bootstrap resampling procedure (1,000 resampling of 500 observations each) with the *boot* package (Canty & Ripley 2015).

This univariate trait-by-trait analysis was complemented with a multivariate analysis. We include these analyses because no trait is truly independent from all others, and it provides an evaluation of joint responses to selection, acknowledging that the evolution of one trait may affect and be affected by the evolution of the others. Furthermore, the best way to measure the evolution of a syndrome, i.e. of multiple traits, is to account for the correlated nature of phenotypic traits (and the modification of these correlations due to selection). Specifically, we performed a Multiple Factor Analysis (MFA) using the *FactoMineR* package (Le et al. 2008). MFA is an extension of a principal component analysis (PCA) where variables can be grouped. For the analysis, we grouped traits by life-history stage. Similarly to a PCA, MFA provides an easy visualization of phenotypic similarities and differences as well as contributions of individual traits to these similarities and differences. Two independent and similar MFAs were conducted for female body mass and age at first reproduction selection schemes. For each selection scheme, each phenotyped trait was averaged over line and diet and included in a global table (except for hatching rate, because no diet was applied prior to hatching). The seven phenotyped traits were gathered into four groups: development (including larval survival rate, development time and age at first reproduction), body mass (including only female body mass), fecundity (including only the number of eggs laid) and survival (including age at death of males and females). Line, diet type, selection type and the interaction between diet type and selection type were added as supplementary qualitative variables and used to interpret the results of the MFA. Dimensions were retained for interpretation when their associated eigenvalue was greater than one. Relationships between variables and between individuals were investigated graphically and using dimension descriptors. Additional graphical representations of trait values and correlations matrices were performed for illustrative purposes, but insufficient replication and high correlations between some trait combinations precluded thorough statistical testing of differences in correlation matrices. All figures were plotted using the *ggplot2* package (Wickham 2016).

## Results

### Direct responses to experimental selection

Female body mass was heritable, with a mean ± s.e. of a broad-sense heritability, H^2^ = 0.46 ± 0.13, and the age at first reproduction was also heritable, with a mean broad-sense heritability H^2^ = 0.19 ± 0.08.

Both selection experiments led to phenotypic changes. Female body mass evolved in both directions during the course of the divergent selection experiment (Figure 1A). At generation 9, female body mass increased by 12% on average in the heavy lines and decreased by 4% in the light lines compared to the control lines (Figure 1B). Age at first reproduction evolved to be significantly earlier in response to selection (χ^2^_2_ = 15.650, p <0.001). At generation 10, age at first reproduction of the replicate lines were between three and 13 days earlier than that of control lines, representing development times that were 29% to 54% faster. These responses to selection led to estimates of narrow-sense heritabilities of h^2^ = 0.11 ± 0.01 for heavy lines and h^2^ = 0.04 ± 0.01 for light lines during selection on female body mass, and h^2^ = 0.10 ± 0.007 for fast lines during selection on the age at first reproduction.

**Figure 1:**
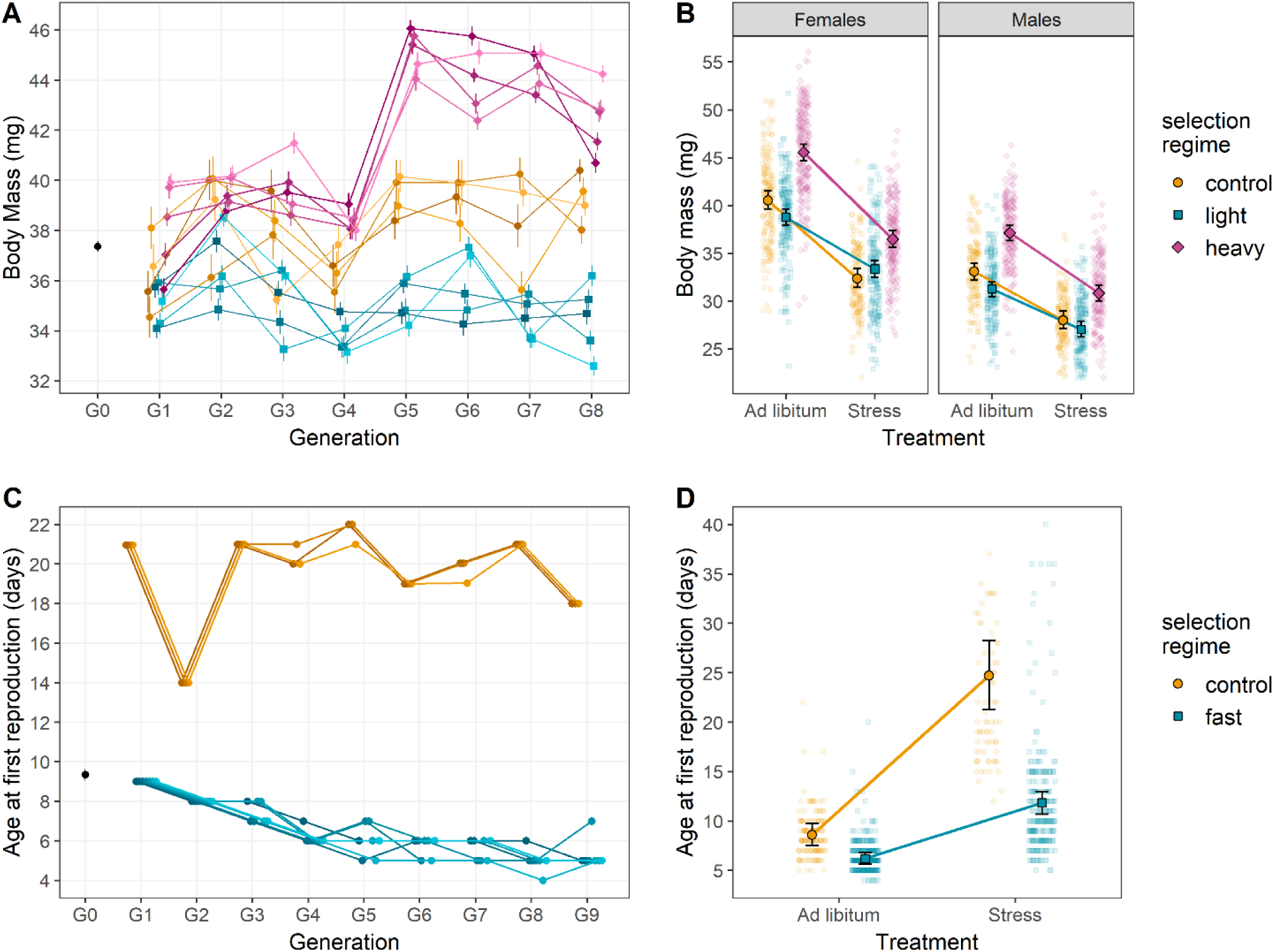
Responses to selection on female body mass and age at first reproduction. (A) Evolution of female body mass for our 14 experimental lines throughout divergent selection on this trait. Heavy, control and light experimental lines are represented in purple, orange and blue, respectively. (B) Body mass of males and females of the G9 generation of female body mass selection. Body masses were measured in either *ad libitum* or stressful nutritional environments. The color code is similar to Fig. 1A. (C) Evolution of female age at first reproduction for our 10 experimental lines throughout directional selection on this trait. Control and selected (fast) lines are represented in orange and blue, respectively. (D) Age at first reproduction for females of the G10 generation of selection on the same trait. Age at first reproduction was measured in either an ad libitum or a stressful nutritional environment. The color code is similar to Fig. 1C. Error bars represent standard errors between replicates.

### Correlated responses to experimental selection

Experimental selection on female body mass did not trigger significant correlated response for most other investigated traits. Hatching rates did not differ among selection types (LRT: χ^2^_2_ = 3.510, p = 0.173; Figure 2A). Similarly, larval survival rate was not influenced by selection on female body mass (LRT: χ^2^_2_ = 0.30, p = 0.86; Figure 2B). Selection on female body mass altered neither development time (LRT: χ^2^_2_ = −1.858, p = 0.395; Figure 2C), nor age at first reproduction (Figure 3A; LRT: χ^2^_1_ = −0.220, p = 0.896). Fecundity did not shift overall with selection on female body mass (Figure 4A; LRT: χ^2^_2_ = 1.35, p = 0.51), but it was influenced by an interaction between selection and diet (see below). While survival differed among sexes, with higher survival in females (LRT: χ^2^_1_ = 107.88, p < 0.0001), it was not influenced by selection on female body mass in females (Figure 4B; LRT: χ^2^_2_ = 1.87, p = 0.39), or in males (LRT: χ^2^_2_ = 1.21, p = 0.27). However, even though the truncation selection was only imposed on females, male mass displayed a correlated response to selection on female body mass with an average increase of 12% in the heavy lines and a decrease of 6% in light lines compared to the control lines in generation 9 (Figure 1B). Additionally, heavy lines seemed to have a higher larval survival rate than light lines (in the *ad libitum* treatment only: multiple comparison of means: Z = 2.275, p = 0.047).

**Figure 2:**
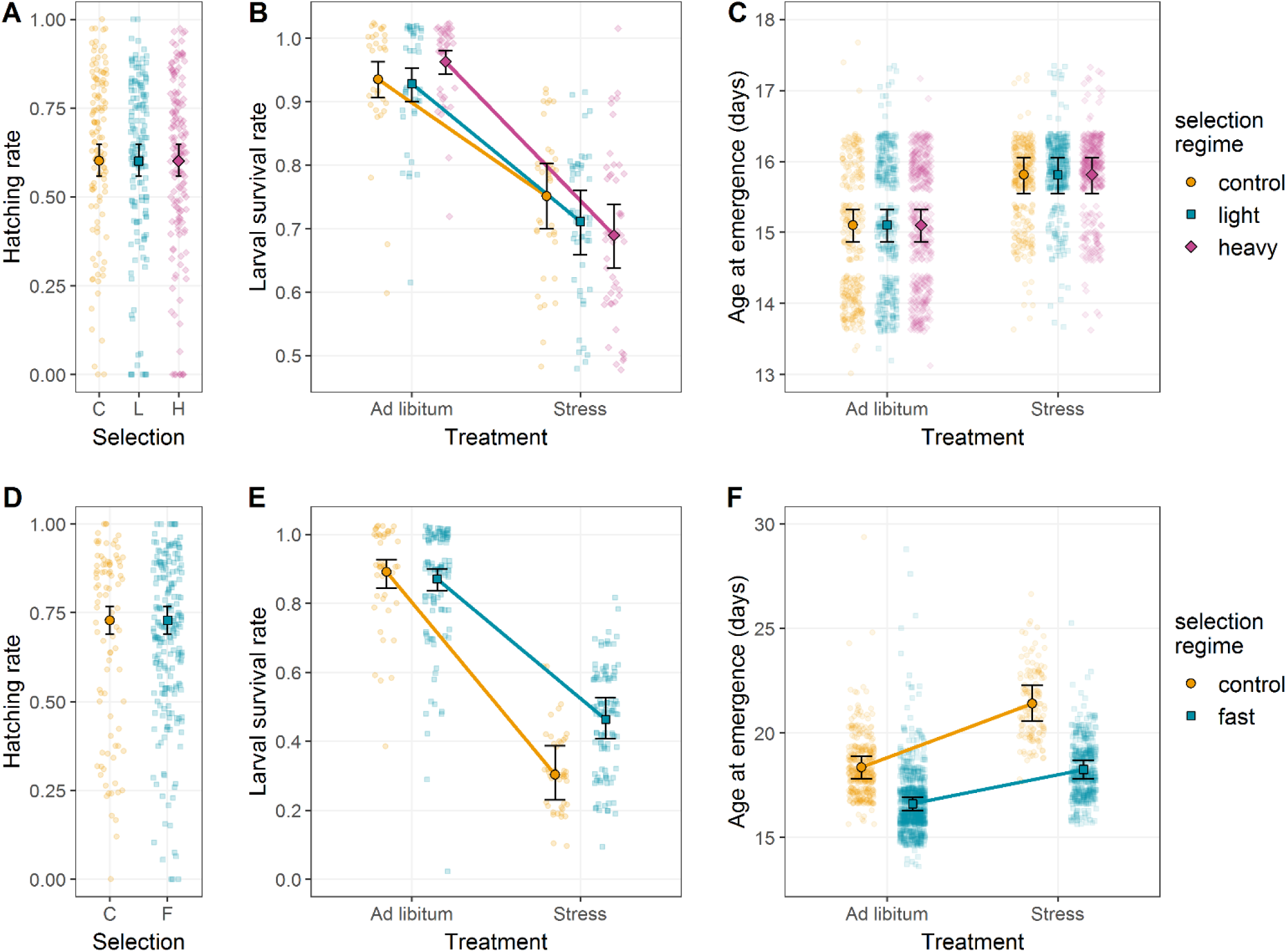
Juvenile phenotypic responses to selection on female body mass and age at first reproduction. Juvenile responses to female body mass selection on (A) hatching rate, (B) larval survival rate and (C) development time. While hatching rate and developmental time were not affected by the selection regime, larval survival rate depended on the interaction between selection regime and developmental environment. In addition, developmental environment had a significant influence on development time. Juvenile responses to age at first reproduction selection on (D) hatching rate, (E) larval survival rate and (F) development time. Hatching rate was not influenced by selection regime, but both larval survival rate and development time were significantly affected by the interaction between selection regime and diet. Color code is similar to Figure 1. Error bars represent standard errors between replicates.

**Figure 3:**
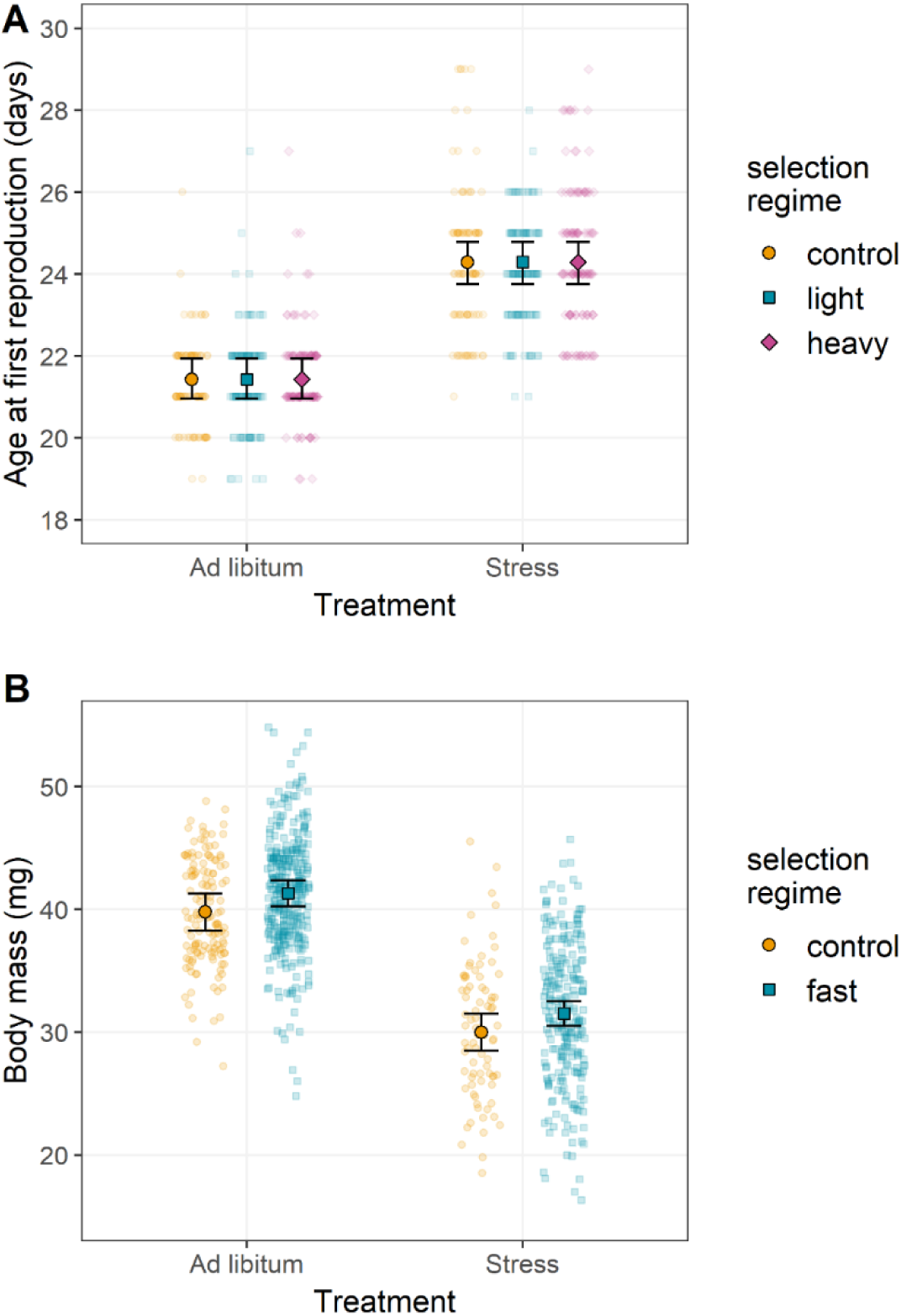
Reciprocal phenotypic responses to selection on female body mass and age at first reproduction. (A) Age at first reproduction of G9 individuals from the selection on female body mass. Diet but not selection regime significantly affected age at first reproduction. (B) Female body mass of G10 individuals from the selection on age at first reproduction. Female body mass was significantly influenced by diet and marginally by selection regime. Color code is similar to Figure 1. Error bars represent standard errors between replicates.

**Figure 4:**
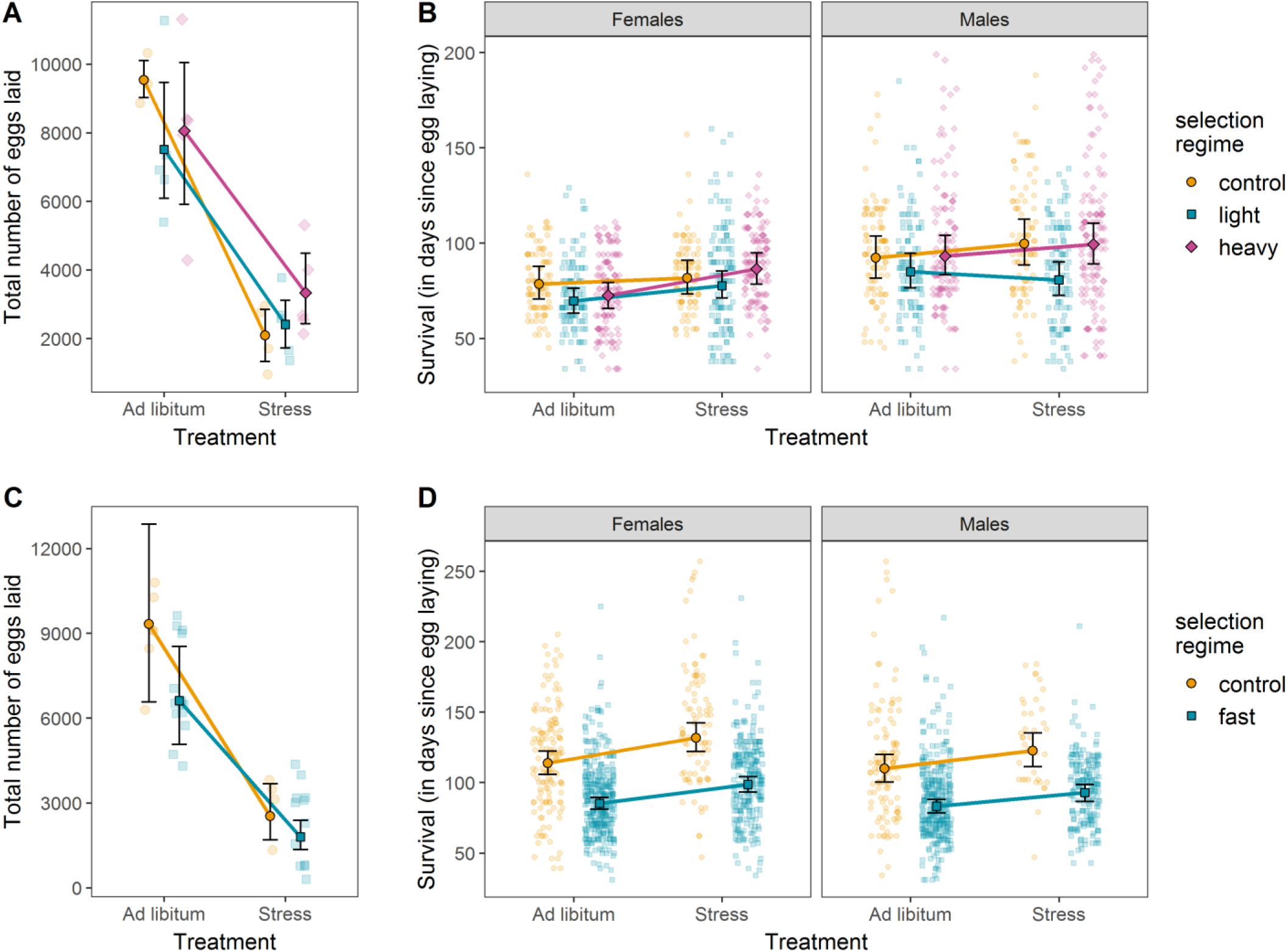
Adult phenotypic responses to selection on female body mass and age at first reproduction. (A) Fecundity of G9 females from experimental lines of the female body mass selection. (B) Female and male survival of G9 individuals of the female body mass selection. (C) Fecundity of G10 females from lines of the age at first reproduction selection. (B) Female and male survival of G10 individuals of the age at first reproduction selection. Color code is similar to Figure 1. Error bars represent standard errors between replicates.

In contrast, experimental selection on age at first reproduction triggered various correlated phenotypic responses, with the exception of hatching rates (LRT: χ^2^_1_ =0.176, p = 0.675; Figure 2D) and larval survival (approx. 90% survival for all lines; Z = −0.777, p = 0.857). Development time, fecundity and survival all responded to selection for fast (i.e. earlier) age at first reproduction, with adults emerging earlier (LRT: χ^2^_1_ = 19.56, p < 0.001; Figure 2F), fecundity decreasing in selected females (LRT: χ^2^_1_ = 4.110, p = 0.043; Figure 4C), and control lines surviving longer than lines selected to reproduce early (Z = 7.280, p < 0.001; Figure 4D). Notably, in contrast to selection on female body mass, which did not affect age at first reproduction, selection on age at first reproduction tended to increase female body mass (best AIC, but LRT only marginally significant: χ^2^_1_ = 2.933, p = 0.087) when provided food *ad libitum* (Wilcoxon rank sum test with continuity correction: p < 0.001; Figure 3B).

### Multivariate response to selection

The Multiple Factor Analyses (MFA) helped visualize the effects of selection (whether targeting female body mass or age at first reproduction) and environmental conditions on trait correlations. We here focus on the effect of selection and will treat the effect of diet in the next section. For both selection experiments, two MFA dimensions had eigenvalues greater than one, hence explaining more variance than any given phenotypic trait alone.

In the female body mass experiment, MFA dimension 1 explained a large proportion of the total variance (58.2%). Traits that contributed to the first dimension were larval survival rate, female body mass, number of eggs laid, development time and age at first reproduction (all correlation coefficients |ρ| > 0.78, all p-values < 0.001; Figure 5A). Dimension 1 separates lines with large females, producing many larvae with short development time and high survival rates (right hand side) from lines with smaller females producing fewer, low-quality larvae. Dimension 2 explained 22.3% of total variance and was mainly due to variance in adult survival (Figure 5A). Dimension 1 clearly separated diet groups (r^2^_diet_ = 0.927, p-value < 0.001), a point that will be detailed in the next paragraph, with selection treatment interacting with diet group (r^2^_interaction_ = 0.965, p-value < 0.001). Dimension 2 correlated weakly with selection treatments (r^2^_selection_ = 0.237, p = 0.034) and strongly with replicate lines (r^2^_line_ = 0.830, p = 0.002).

**Figure 5:**
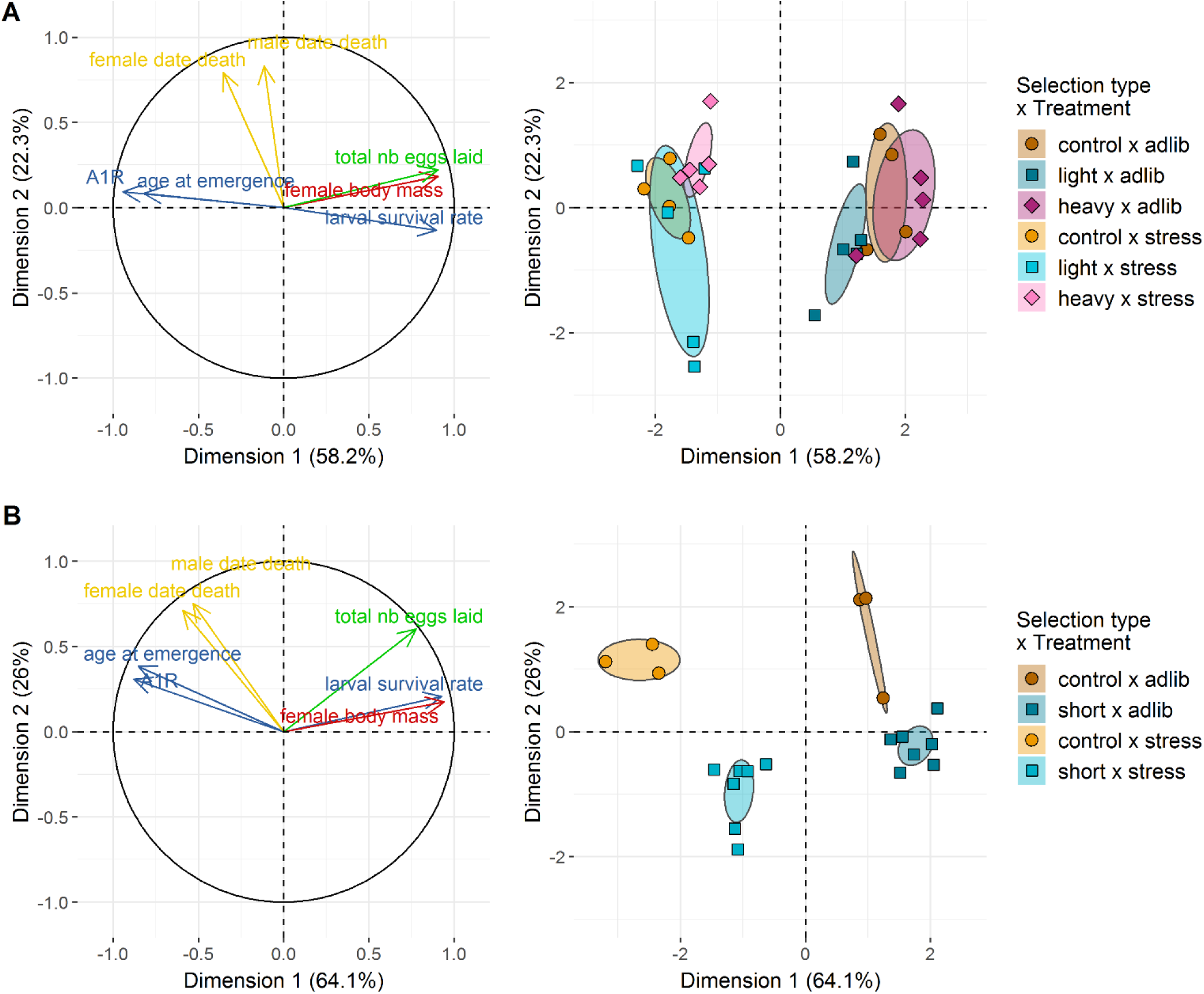
Organismal-level phenotypic responses to selection and diet. (A) right: Projection of phenotypic traits into the first two dimensions of the Multiple Factor Analysis (MFA) on G9 individuals of the female body mass selection regime. A1R stands for age at first reproduction. (A) left: Projection of replicate experimental lines from selection on female body mass into the first two dimensions of the same MFA. Diet is a major factor separating global phenotypes while selection regime is a more modest one. (B) right: Projection of phenotypic traits into the first two dimensions of the MFA on G10 individuals of the age at first reproduction selection. When comparing (A) right and (B) right, it is worth noting that the relationships between phenotypic traits are mostly similar on Dimension 1 (i.e., related to diet treatment) but are more dissimilar on Dimension 2 (i.e., related to selection regime). (B) left: Projection of replicate lines from selection on age at first reproduction into the first two dimensions of the same MFA. Both the diet and selection regime are participating in the clear-cut clustering of individual lines’ phenotypes, contrary to (B) where diet and the interaction between diet and selection regime are the main clustering factors.

Overall this MFA showed that selection on female body mass only weakly influenced the distribution of trait values, with lines belonging to different selection treatments not completely separated in the MFA plane. The phenotypic correlation matrices (Supplementary Figure 4) corroborate this conclusion. Some selected lines had trait combinations close to those of control lines while other selected lines differed from the control lines. The lines that were the most distinct from control lines, tended to have either greater (heavy lines) or smaller (light lines) coordinates on MFA Dimension 2 compared to control lines, an indication that selection on female body mass might have altered other traits in these lines. Radar plots provide a visual confirmation that global phenotypic variation due to selection on body mass was low, except for body mass itself (Figure 6, top row), with marginal effects on adult survival in some selected lines.

**Figure 6:**
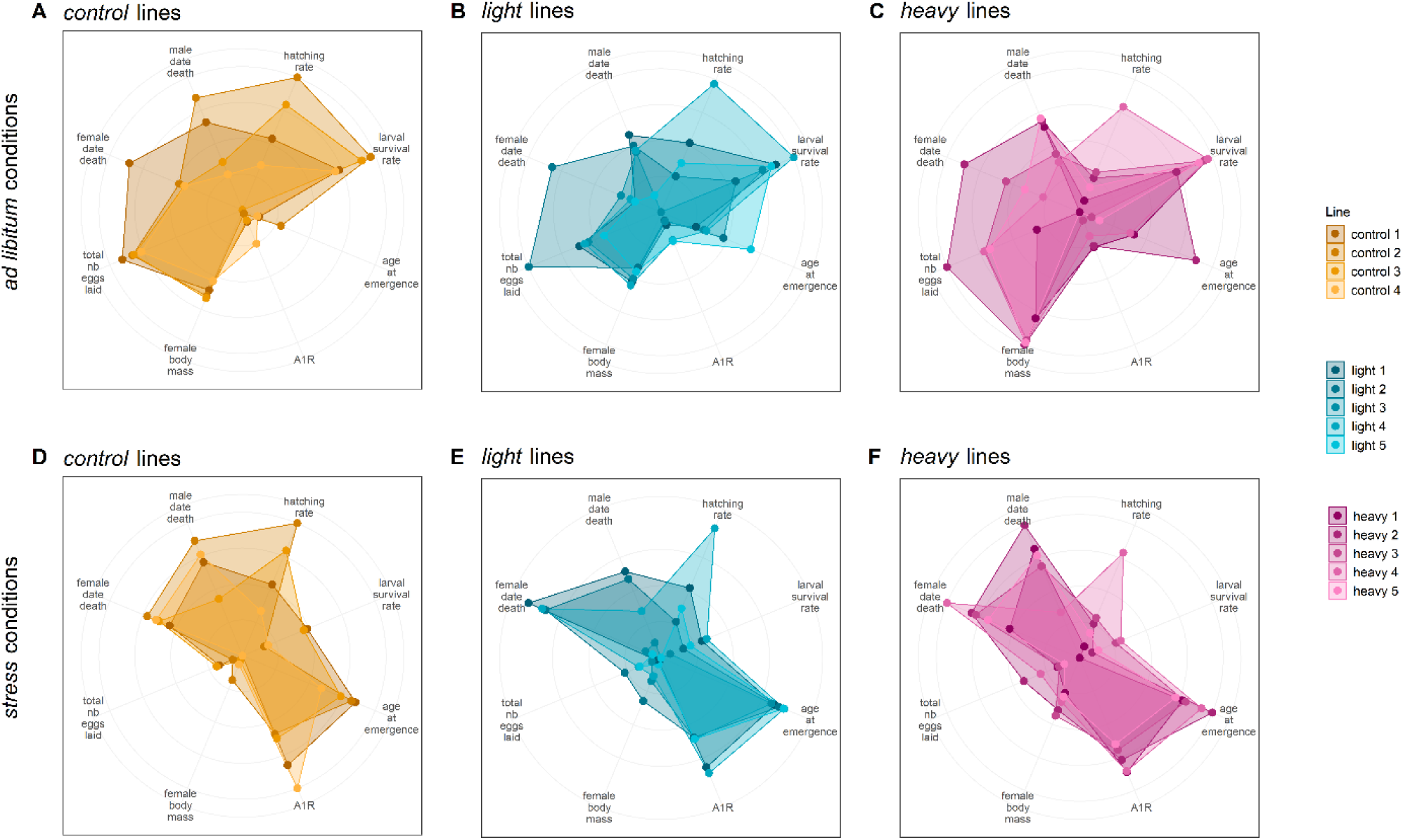
Multi-trait phenotypes of experimental lines of the female body mass selection scheme. (A-C) Radar plots of G9 control, light and heavy lines phenotyped under ad libitum conditions for eight traits. The lines display a strong phenotypic response on the selected trait (female body mass), while the rest of their phenotype remains similar. A1R stands for age at first reproduction. (D-F) Radar plots of G9 control, light and heavy lines phenotyped under stressful conditions for eight traits. The lines display a similar phenotypic response, which is strongly dissimilar to the phenotypic response obtained under ad libitum conditions. Color code is similar to Figure 1.

In the age at first reproduction experiment, dimensions 1 and 2 explained 64.1% and 26% of the variance, respectively. The contributions of traits to both dimensions were relatively similar to the body mass selection experiment, with however looser correlations between dimension 2 and adult survival rates, and between dimension 1 and age at first emergence (Figure 5B). Here, male and female survival were also significantly negatively correlated with the MFA dimension 1 (ρ = −0.53, p = 0.016 and ρ = −0.59, p = 0.006, respectively; Figure 5B). In contrast to selection on female body mass, both dimensions clearly separated both diet and selection groups (Figure 5B). Dimension 1 strongly correlated with diet (r^2^_diet_ = 0.857, p-value < 0.001) and the interaction between diet and selection (r^2^_interaction_ = 0.974, p-value < 0.001), Dimension 2 correlated strongly with both selection treatment (r^2^_selection_ = 0.717, p < 0.001) and line (r^2^_line_ = 0.813, p = 0.011). Lines tended to group by diet and selection treatments revealing a strong effect of selection on the distribution of the other traits. Radar plots confirm that the global phenotypic variation due to selection on age at first reproduction was high (Figure 7, top row). Correlation matrices showed that selection on the age at first reproduction modified phenotypic correlations, with a noticeable relaxation of correlations between traits in the selected lines (Supplementary Figure 5, top row). These multivariate analyses show that correlations between traits has been profoundly been modified by selection on age at first reproduction, with some traits positively responding to the treatment (e.g., adult survival rates) and others not.

**Figure 7:**
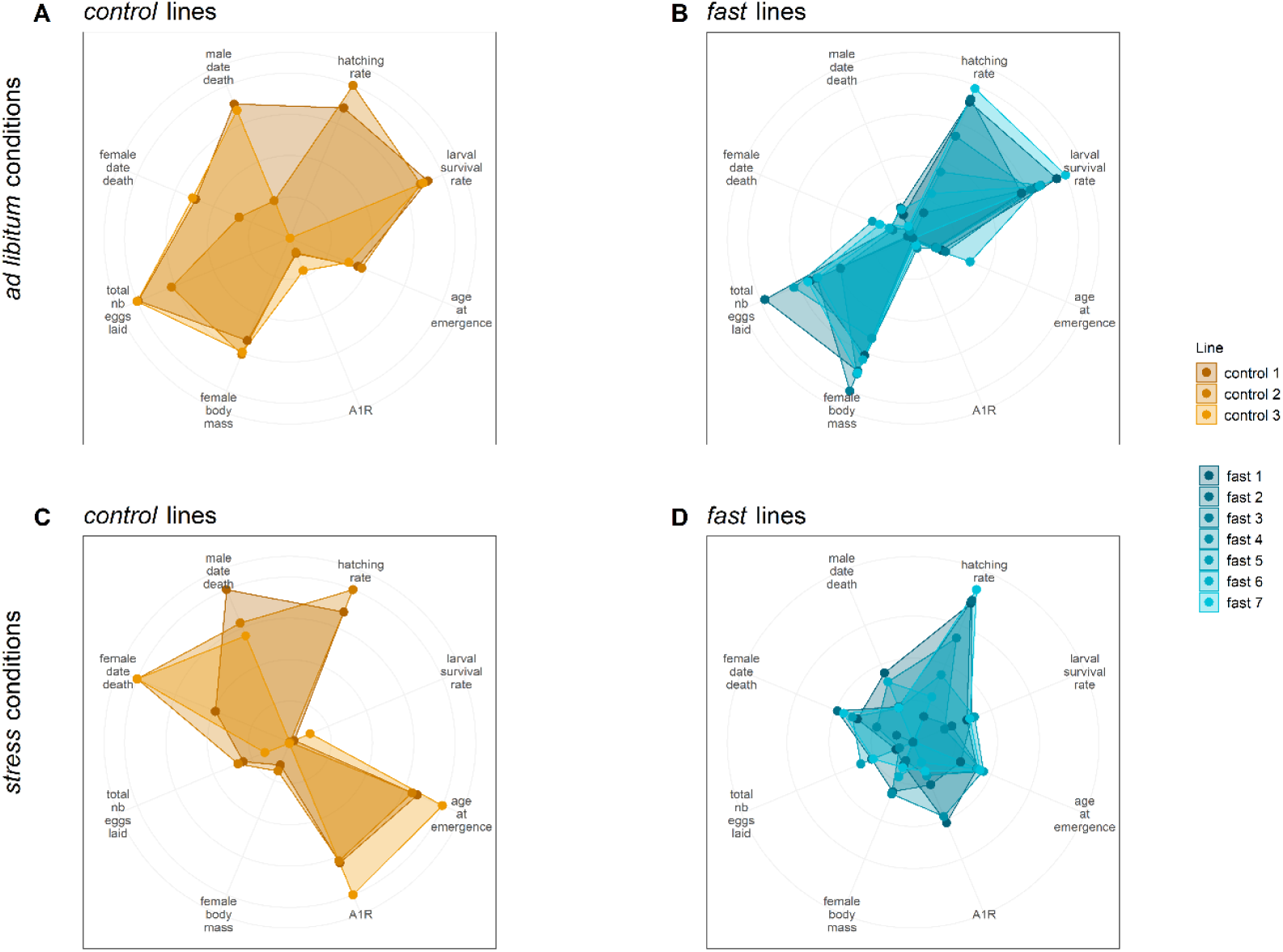
Multi-trait phenotypes of experimental lines of the age at first reproduction selection scheme. Radar plots of G10 control and fast lines phenotyped under ad libitum conditions (A-B, respectively) and stressful conditions (C-D, respectively) for eight traits. Each line displays a distinctive phenotypic response to selection × environment conditions. In contrast to the female body mass selection, selection on age at first reproduction modified most investigated traits. Color code is similar to Figure 1.

### Environmental context and trait expression

In both selection experiments, we found that the environmental context, here expressed in the form of abundant versus reduced resources during the final phenotyping step, played a major role in explaining the phenotypic syndromes, often in interaction with selection.

After experimental selection on female body mass, food stress reduced the mass of females and males as expected, with a significant interaction between the diet type and the direction of the selection (Figure 1A; females: χ^2^_2_ = 36.337, p <0.001; males: χ^2^_2_ = 15.288, p <0.001). The difference in mass between heavy and light lines was always more pronounced when fed *ad libitum* than under stress for both sexes (multiple comparisons of means: females: Z = 5.875, p < 0.001; males: Z = 3.923, p < 0.001). Larval survival was also influenced by the interaction between selection type and diet type (Figure 2B; χ^2^_2_ = 7.397, p = 0.025). Heavy lines survived at a higher rate than light lines in the *ad libitum* but not in the stressful diet (multiple comparison of means: Z = 2.275, p = 0.047). On the other hand, heavy lines survived at a lower rate than control lines in the stress but not in the *ad libitum* diet (Z = −2.364, p = 0.047). The difference between control and light lines did not depend on diet (Z = −0.264, p = 0.792). Adults emerged earlier when they had abundant food than when under food stress (Figure 2C; LRT: χ^2^_2_ = −16.683, p < 0.001), but no interaction with selection was present for this trait (LRT: χ^2^_2_ = −0.344, p = 0.842). While selection on female body mass did not alter the age at first reproduction (Figure 3A), diet did (LRT: χ^2^_1_ = 59.23, p < 0.001), with reproduction delayed for all lines under food stress, whether selected to be heavy or light (+2.86 days on average; no selection × diet interaction: χ^2^_2_ = −0.145, p = 0.930). Fecundity was influenced by an interaction between selection on female body mass and diet (Figure 4A; LRT: χ^2^_2_ = 4.730, p = 0.030). Specifically, control lines suffered a more severe drop in fecundity when food was limited than selected lines (control vs. heavy lines: Z = 3.038, p = 0.007; control vs. light lines: Z = 1.855, p = 0.064). The diet type affected heavy and light lines in a similar manner (Z = −1.255, p = 0.210). Survival was affected by a 3-way interaction between sex, selection and diet (Figure 4B; LRT: χ^2^_2_ = 6.378, p = 0.041). We thus analyzed the survival results according to sex. For females, selection type interacted with diet (LRT: χ^2^_2_ = 6.378, p = 0.017). This interaction was mostly driven by the increase in female survival in heavy lines under stress when compared to control females (Z = 2.863, p = 0. 012). For males, the interaction between the selection type and the diet type was also statistically significant (LRT: χ^2^_2_ = 8.180, p = 0.020), mostly driven by the drop in survival in males from light lines under stress in comparison to heavy and control lines (Figure 4A).

After experimental selection on the age at first reproduction, the earlier female reproduction we observed was magnified under the stressful diet (selection × diet interaction: χ^2^_2_ = 35.567, p <0.001), with fast lines laying their first eggs on average three days earlier than control lines under the *ad libitum* diet (fast lines = 6 days, control lines = 9 days), and as much as 13 days earlier in the stress diet (fast lines = 12 days, control lines = 25 days, Figure 1D). Food stress reduced larval survival overall (all comparisons: p < 0.001), with greater reductions in the fast lines than in the control lines (46% and 30% survival for fast and control lines, respectively; selection × diet interaction: χ^2^_1_ = 13.434, p < 0.001; Figure 2E). Adults emerged earlier when they had abundant food (Figure 2F; LRT: χ^2^_1_ = 91.88, p < 0.001). Development time was also influenced by an interaction between selection and diet (LRT: χ^2^_1_ = −5.709, p = 0.017), with the control lines under food limitation taking the longest to emerge. Fecundity also responded to diet, with food stress strongly reducing fecundity (LRT: χ^2^_1_ = 51.200, p < 0.001), but without interacting with selection (LRT: χ^2^_1_ = 0.188, p = 0.665). Finally, survival increased with food stress (LRT: χ^2^_1_ = 14.14, p < 0.001, Figure 4D), without any interaction between other significant factors (such as sex and selection; all p > 0.12).

Multivariate analyses of the responses to selection highlighted a strong effect of the diet type on phenotypes. As mentioned above, for both experimental selection schemes, MFA dimension 1 significantly correlated with diet and with the interaction between diet and selection (female body mass: r^2^_diet_ = 0.927 and r^2^_interaction_ = 0.965, both p-values < 0.001; age at first reproduction: r^2^_diet_ = 0.857 and r^2^_interaction_ = 0.974, both p-values < 0.001); Figure 5). In both experiments, within each selection treatment, stress and *ad libitum* lines tended to separate along the direction given by female body mass quasi-orthogonally to that of adult survival rates. Stressed lines therefore had trait combinations dominated by small females, few low-quality larvae with long development and late reproduction, with no major effect on adult survival. This MFA also revealed a large interaction effect with selection corresponding to different responses to diet according to the selection type. For instance, in Figure 5B heavy lines are distinct from control and light lines under food stress, while light lines are distinct from control and heavy lines when fed *ad libitum*. Diet and diet × selection effects were most obvious when comparing *ad libitum* and stress phenotypes in radar plot illustrations and correlation matrices (Figures 6-8, Supplementary Figures 4 and 5, top vs. bottom rows).

**Figure 8:**
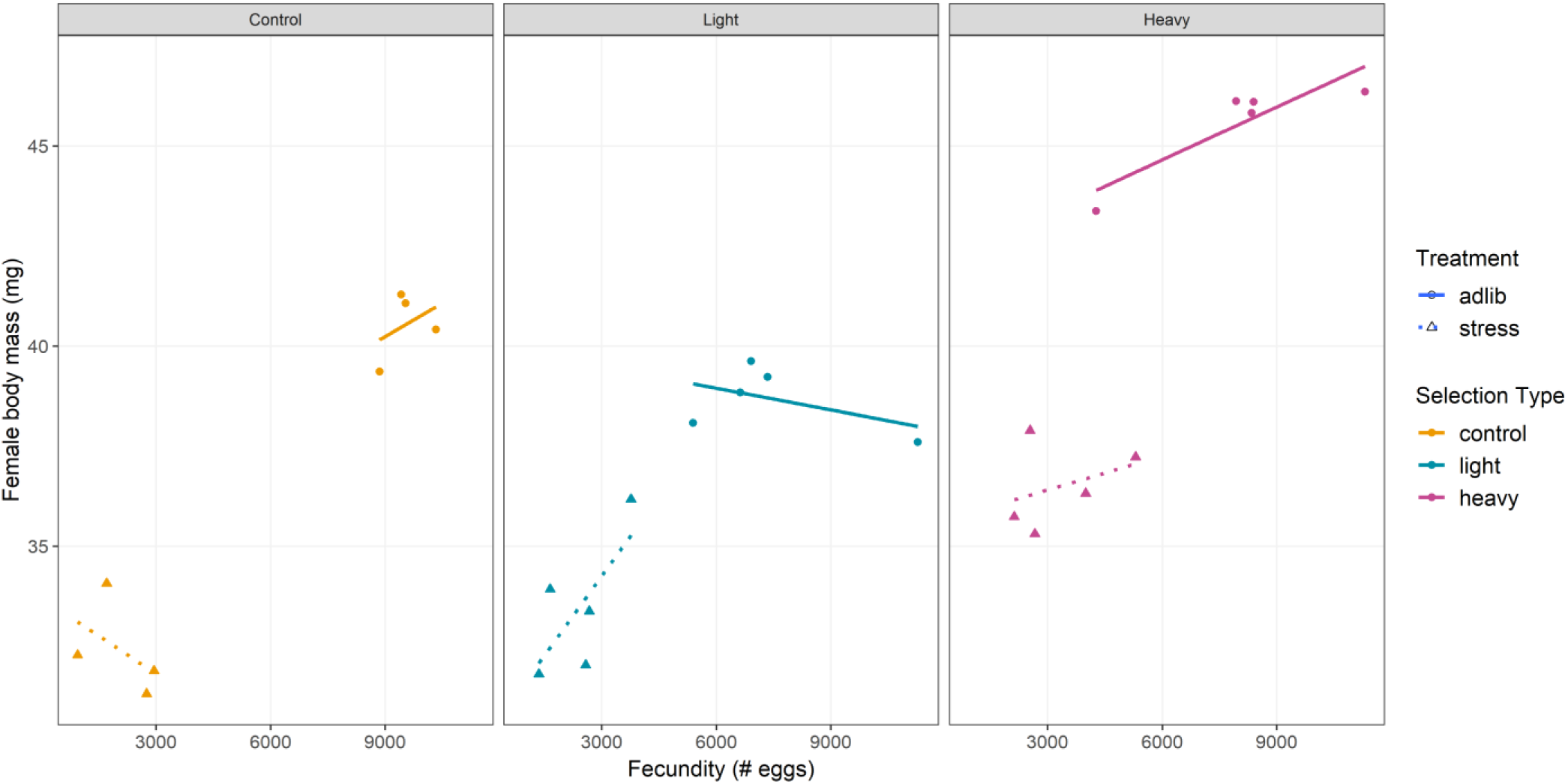
Relationship between reproductive and growth traits of experimental lines of the female body mass selection scheme. Each point refers to a replicate line in a given experimental condition. Dots and triangles indicate lines in ad libitum and stressful conditions, respectively. Solid and dashed lines represent linear regression for ad libitum and stressful conditions, respectively. Color code is similar to Figure 1.

## Discussion

We investigated the mechanisms underlying the emergence of the invasion syndrome occurring in invasive populations of *H. axyridis* using two distinct artificial selection experiments performed on two traits involved in this syndrome. This suite of experiments enabled us to assess whether (i) the selected traits evolved, (ii) other phenotypic traits evolved in concert, (iii) the new phenotypic syndromes resembled the invasion syndrome found in natural populations of *H. axyridis*, and (iv) environmental conditions impacted the expression of the evolved syndromes.

### Rapid evolution of traits under direct selection

The two focal traits, female body mass and age at first reproduction, were heritable (H^2^ = 0.46 and 0.19 for female body mass and age at first reproduction, respectively) and responded to selection. In less than ten generations, female body mass increased by 12% in the heavy lines and decreased 4% in the light lines compared to the control lines and the age at first reproduction declined by 33% in the fast lines compared to the control lines in *ad libitum* conditions. In contrast, control lines displayed no evolution of age at first reproduction and only a slight increase of female body mass compared to the initial generation, indicating no ongoing evolutionary response to rearing conditions. These results confirm several studies showing that in many species body mass or size and age at first reproduction are heritable and can rapidly evolve in artificial selection experiments (e.g. Teuschl et al. 2007, Miyatake 1997), as well as in natura especially in the context of biological invasion (e.g. Reznick et al. 1990, Huey et al. 2000, Diamantidis et al. 2009, Kingsolver et al. 2012). The fact that single traits linked to growth or reproduction can quickly respond to selection was expected, due to the phenotypic changes that we observed for this type of traits between native and invasive populations of *H. axyridis* (Supplementary Figure 1).

### The evolution of a phenotypic syndrome highly depends on the selected trait

The main objective of our work was to assess whether other phenotypic traits evolved in response to selection on body mass or age at first reproduction, and to evaluate the extent to which such correlated shifts mimicked the invasion syndrome. Life-history theory has given keys to understand how different traits could interact to produce different life-history strategies (Stearns 1992). Faced with new selection pressures, it is quite common that a suite of traits will evolve rather than a single trait in isolation, whether in a context of experimental evolution or in natura (Reznick et al. 1990, Reznick & Ghalambor 2001, Teuschl et al. 2007). Here we found that the number of traits showing a correlated response is highly variable depending on the trait undergoing direct selection.

The impact of selection on female body mass on the other studied traits was weak when measured in *ad libitum* conditions. Male body mass showed a correlated response to selection, suggesting that genes determining weight are mostly on autosomal chromosomes. Larval survival rate also displayed a slight correlated response, with larvae from heavy lines surviving better than those from light lines. Finally, fecundity tended to decrease for both heavy and light lines compared to control lines. Overall, while selection on female body mass was highly efficient in both directions, it did not lead to the emergence of a complex phenotypic syndrome such as the one observed in invasive populations of *H. axyridis*. At first glance, this result seems contradictory with the fact that most life-history traits often covary with size (Woodward et al. 2005). However, observing weak correlated responses of other traits to direct selection on a single focal trait is not uncommon in experimental selection assays (Spitze et al. 1991). A first potential explanation is related to the relatively short duration of our artificial selection experiments, which led to a female body mass increase of 12%. In comparison, Malerba et al. (2018) used 280 generations of artificial selection to evolve a 10-fold difference in mean body size between small- and large-selected lineages of the phytoplankton *Dunaliella tertiolecta*. In the Malbera et al. (2018) experiment, the intense selection on the size resulted in a shift in several traits involved in the ability of this species to cope with resource fluctuations. It is possible that *H. axyridis* would have exhibited such shifts over time. A second potential explanation relies on the fact that presumed trade-offs underlying the evolution of adaptive life histories often become visible only in stressful environments (Schluter et al. 1991). Finally, although metabolic theory explains the covariation between size and demography across species, this pattern does not seem to hold within a species and the link between size and other life-history traits within species could be less direct than commonly thought (Malerba et al. 2019).

In contrast, artificial selection on age at first reproduction resulted in a strong correlated response in most of the other traits that were studied. We observed a strong shift in four of the six traits (development time, body mass, fecundity and survival). Only the earliest developmental traits did not show a correlated response (i.e., hatching rate and larval survival). In *ad libitum* phenotyping conditions, individuals from fast lines developed faster and died earlier than control lines, and females from fast lines were bigger and laid fewer eggs than control lines. Thus, selection on a single trait (here age at first reproduction) resulted in the evolution of a clearly distinct multi-trait phenotypic syndrome.

Overall, our results suggest that all traits are not equivalent and symmetric in building a genetic response to selection. It seems that selection on age at first reproduction drove the evolution of female body mass, but that the reverse was not true. Due to technical constraints, the two experiments had to be performed separately, and thus it is not impossible that the observed differences in the response to selection are due to other differences in the experiments. Most notably, the experiments started with different base populations. The base population for selection on female body mass experienced eight generations in the laboratory environment before the start of artificial selection process, while the base population for selection on age at first reproduction experienced only two generations in the lab. Since base populations were not lab adapted at the start, this may mean that during selection on age at first reproduction, the populations, also could have been experiencing selection to the laboratory environment. To test this, we compared the G0 founding population with control lineages at the end of selection (G10). Both univariate and multivariate analyses indicate that most traits evolved only slightly or not at all between G0 to G10 in the absence of selection on age at first reproduction (results not shown, R code and data available in the online supplementary material). We tested a potential impact of differences between experiments by re-running all analyses using the difference between observed trait values and the means of controls lines rather than the raw trait data. These analyses yielded the same results as raw-value analyses, indicating a lack of influence of different experimental settings on the conclusion (results not shown, R code and data available in the online supplementary material, https://data.inra.fr/dataset.xhtml?persistentId=doi:10.15454/V9XCA2). In addition to the evolution of trait values, the relationships between the phenotypic traits were modified by the selection experiments. Multivariate analyses confirmed that selection for a faster age at first reproduction modified the phenotype more profoundly than selection for female body mass. Interestingly, we found that selection on the age at first reproduction (and to a lower extent on the body mass), resulted in a global relaxation of phenotypic correlations, and this was independent of the environmental context (i.e. the diet type during the phenotyping step). The correlated responses varied highly depending on the trait under direct selection. This indicates that selective pressures, even when transient like in our experiments, may have profound impacts on additive genetic variance-covariance matrices (G-matrices) and therefore on future evolutionary trajectories (Steppan et al. 2002; Arnold et al. 2008, Blows & Mcguigan 2015). Overall, our results show that the phenotypic matrices are far from rigid and can evolve rapidly due to both selective and environmental contexts, indicating that selection on different traits will have different consequences for the relationships between traits. Rapid evolution of correlated traits following introduction into a novel environment might be common, as illustrated by correlated morphological and behavioral traits evolution following colonization of a new habitat in the isopod Asellus aquaticus (Eroukhmanoff & Svensson 2011; Karlsson Green et al. 2016).

### Experimental selection to study invasion syndromes

A key question of the present study was whether selection on one trait of the invasion syndrome observed in invasive populations of *H. axyridis* could lead to inducing the whole or at least a part of the syndrome in the laboratory. We have compiled the correlated responses observed in *ad libitum* phenotyping conditions for the two artificial selection experiments as well as the invasion syndrome observed *in natura* in Table 1. Regarding female body mass, artificial selection has clearly resulted in few correlated responses and hence poorly reproduced the invasion syndrome. In contrast, for age at first reproduction, artificial selection resulted in more widespread phenotypic changes, which nevertheless corresponded only partly to the invasion syndrome. Similar to invasive populations, artificial selection on age at first reproduction triggered faster larval development, earlier age at reproduction and heavier body mass. However, we observed several trade-offs that were not found in invasive populations: individuals from fast lines exhibited a shorter lifespan and had a lower egg production than individuals from control lines.

The discrepancy between phenotypes of natural invasive populations and experimentally selected lines has several potential explanations. First selection in natura was longer than 10 generations of artificial section in the lab. Second selective pressures are likely more complex during the course of the invasion as compared to the laboratory, probably involving a large set of biotic factors and biotic interactions (Reznick and Ghalambor 2001, Mitchell et al. 2006). Moreover selection pressures *in natura* can be variable through time (Sakai et al. 2001), which would greatly alter the phenotypic outcome of selection. Finally, the successful settlement of introduced propagules followed by demographic expansion leading to invasion is as a trial and error process (Laugier et al. 2016) from which we only see the winners (McKinney and Lockwood 1999). In the case of *H. axyridis*, multiple introductions for biological control occurred prior to invasion, and the groups that did finally invade may have been superior in some way that enabled them to escape the trade-offs we observed between time to first reproduction, lifespan and fecundity in our selection experiment. It remains unknown whether the invasion and its associated syndrome were successful due to such lottery effects or because of the purging of deleterious during the multiple introductions, creating individuals with overall higher fitness (Facon et al. 2011).

### Environmental features strongly influence the expression of phenotypic syndromes

Our experiments revealed a large impact of diet on the evolutionary shifts observed for all measured traits. In agreement with the study of Sikkink et al. (2017), we found that the effect of diet was far from being homogeneous and was very dependent on the trait considered and the selection regime. The phenotypic differences between control and selected lines were magnified in stressful conditions for the fast reproduction lines. Differences between heavy lines and light and control lines were exacerbated under stressful diet. Interestingly, control lines displayed a positive relationship between growth and reproductive traits in *ad libitum* conditions and a negative relationship between the same traits in stressful conditions (Figure 8). These observations are reminiscent of the *Daphnia pulex* balance between ‘superfleas’ showing no phenotypic trade-offs in favorable environments, while stressful conditions reveal a cost of acquisition leading to allocation trade-offs (Reznick et al. 2000). Such dependence of trade-offs on environmental conditions is interpreted as revealing a cost of acquisition where ‘high acquisition-low allocation’ phenotypes are favored in situations where resources are abundant and ‘low acquisition-high allocation’ phenotypes are favored in low resources environments (Reznick et al. 2000).

Selection for lighter females seemed to have reversed this pattern, with light lines showing no trade-off between growth and reproduction in stressful phenotyping conditions, while a trade-off appeared in *ad libitum* conditions. Moreover, selection for heavier females might have triggered a ‘high acquisition’ phenotype that displays a lack of genetic variation in resource allocation (Reznick et al. 2000), a pattern particularly marked in stressful conditions (e.g., positive slope for heavy × stress lines in Figure 8). Different directions of selection thus seem to have triggered new responses in the allocation and acquisition of available diet resources. This pattern however deserves a dedicated experimental study to evaluate the influence of past selection on the balance between acquisition and allocation of diet resources.

By supporting that adaptation in one environment may induce benefits and costs in another environment (Dutilleul et al. 2017), our results underline the importance that G × E interactions could have for the studies of biological invasions. Indeed, native and invaded environments are likely to be different (Reznick and Ghalambor 2001) and the invaded environments may themselves undergo substantial changes during the course of the invasion (Sakai et al. 2001). In the case of *H. axyridis*, the agricultural habitats in which *H. axyridis* was introduced as a biocontrol agent are certainly different in terms of nutritional resources and habitat structures from the habitats encountered later during its geographical expansion. Moreover, the eco-evolutionary dynamics of spreading populations may also be constrained due to the balance between selective and stochastic processes (Williams et al. 2019). Overall, our results highlight the need to systematically examine the consequences of experimental evolution in a variety of environments (Amarillo-Suarez et al. 2011).

## Conclusion

The adaptive challenges encountered by a species in its introduced habitat are complex. Manipulative field experiments imposing selection are difficult to impossible to implement, particularly with invasive species. Thus, artificial selection can be employed to mimic what could have happened during the course of invasion, evaluate correlated responses to selection for focal traits and estimate the extent to which those correlations shape the suite of traits specific of the invasive populations (Fuller et al. 2005). Our artificial selection experiments failed, however, to reproduce the complete invasion syndrome we documented in natural populations of *H. axyridis*. This result underlines the limits of using laboratory experiments to study complex evolutionary trajectories in natural conditions. To go further in the evaluation of the common points and discrepancies between artificial selection lines and natural populations, it would be relevant to carry out whole-genome scans to compare the genomic regions showing signals of selection associated with invasive natural populations to the genomic regions showing signals of selection in our experimental lines. The laboratory experiments we carried out are nonetheless a mandatory step in identifying the traits that are correlated and those that are less so in various selective contexts, helping us understand the evolution of an invasion syndrome. Finally, the strong G × E interactions we observed indicate that trait values adaptive in one environment may no longer be advantageous following environmental change. This could explain, at least partly, why boom-bust dynamics – the rise of a population to outbreak levels, followed by a strong decline – have been frequently described in invasive populations (Lockwood et al. 2013; Strayer et al. 2017).

## Data accessibility

All data and R code are available online at: https://data.inra.fr/dataset.xhtml?persistentId=doi:10.15454/V9XCA2

## Supplementary material

Supplementary material is available online at: https://www.biorxiv.org/content/10.1101/849968v2.supplementary-material

## Acknowledgements

AE and BF acknowledge financial support by the national funder ANR (France) through the European Union program ERA-Net BiodivERsA (project EXOTIC), and the INRA scientific department SPE. AE and JF acknowledge the UMR CBGP from its internal grant support. VR acknowledges financial support by the European Union regional development fund (ERDF), the Conseil Départemental de la Réunion, the Région Réunion, and the French Agropolis Fondation (Labex Agro–Montpellier, E-SPACE project number 1504-004). RAH acknowledges support from USDA AFRI COLO‐2016‐ 09135, USDA NIFA Hatch project 1012868, the French Agropolis Fondation (LabEx Agro–Montpellier) through the AAP ‘International Mobility’ (CfP 2015-02), the French programme investissement d’avenir, and the LabEx CEMEB through the AAP ‘invited scientist 2016’. We are grateful to the SEPA technical platform of the CBGP laboratory for hosting all experiments presented in this study. JF would like to thank Pierre E. Sirius and Ariane N. Véga for their early support. Version 4 of this preprint has been peer-reviewed and recommended by Peer Community in Evolutionary Biology (https://doi.org/10.24072/pci.evolbiol.100096).

## Conflict of interest disclosure

The authors of this preprint declare that they have no financial conflict of interest with the content of this article. RAH, VR, AE and BF are PCI Evol Biol recommenders.

## Notes

https://data.inra.fr/dataset.xhtml?persistentId=doi:10.15454/V9XCA2

### Competing Interest Statement

The authors have declared no competing interest.

### Summary of Updates

Manuscript and figures updated following Peer Community in Evolutionary Biology revision round 2. Datasets and R code made public.

