## Supplementary Material for "How do invasion syndromes evolve? An experimental evolution approach using the ladybird Harmonia axyridis"

RESEARCH ARTICLE

Open Access
Open Data
Open Code
Open Peer-Review

### How do invasion syndromes evolve? An experimental evolution approach using the ladybird Harmonia axyridis

Julien Foucaud[1,*], Ruth A. Hufbauer[1,2], Virginie Ravigné[3], Laure Olazcuaga[1], Anne Loiseau[1], Aurélien Ausset[1], Su Wang[4], Lian-Sheng Zang[5], Nicolas Leménager[1], Ashraf Tayeh[1], Arthur Weyna[1], Pauline Gneux[1], Elise Bonnet[1], Vincent Dreuilhe[1], Bastien Poutout[1], Arnaud Estoup[1], Benoît Facon[6,*]

[1] UMR CBGP (INRA-IRD-CIRAD, Montpellier SupAgro), Campus International de Baillarguet, CS 30 016, 34988 Montferrier / Lez cedex, France

[2] Colorado State Univ, Dept Bioagr Sci & Pest Management, Graduate Degree Program in Ecology, Ft Collins, CO 80523 USA

[3] CIRAD, UMR PVBMT, F-97410 Saint-Pierre, Réunion, France

[4] Beijing Academy of Agriculture and Forestry Sciences, China

[5] Institute of Biological Control, Jilin Agricultural University, Changchun, China

[6] INRA, UMR PVBMT, F-97410 Saint-Pierre, Réunion, France

**Cite as:** Foucaud J, Hufbauer RA, Ravigne V, Olazcuaga L, Loiseau A, Ausset A, Wang S, Zang LS, Leménager N, Tayeh A, Weyna A, Gneux P, Bonnet E, Dreuilhe V, Poutout B, Estoup A, and Facon B. How do invasion syndromes evolve? An experimental evolution approach using the ladybird *Harmonia axyridis*. bioRxiv 849968, ver. 4 peer-reviewed and recommended by PCI Evolutionary Biology (2020).

**Posted:** 23rd April 2020

**Recommender:**
Inês Fragata and Ben Philipps

**Reviewers:**
Two anonymous reviewers

**Correspondence:**



This article has been peer-reviewed and recommended by
*Peer Community in Evolutionary Biology*

### Supplementary Material

**Supplementary Figure 1: Female body mass and female age of first reproduction for invasive and native populations of H. axyridis**

Data were produced during the experiment that led the following publication: Tayeh, A., Hufbauer, R. A., Estoup, A., Ravigné, V., Frachon, L., Facon, B. (2015). Biological invasion and biological control select for different life histories. Nature Communications, 6. 7268, DOI : 10.1038/ncomms8268.

(A) For female body mass, data are from 132 individuals coming from two invasive (n=33 for Santiago in Chile, n=43 for Brookings in USA) and two native (n=28 for Beijing in China, n=28 for Fuchu in Japan) populations. Invasive females are heavier than native ones (Wilcoxon signed-rank test: $W=3147.5$, $p < 0.0001$).

(B) For age at first reproduction, data are from 114 individuals coming from two invasive (n=27 for Santiago in Chile, n=36 for Brookings in USA) and two native (n=25 for Beijing in China, n=26 for Fuchu in Japan) populations. Invasive females initiate reproduction earlier than native ones (Wilcoxon signed-rank test: $W=3147.5$, $p < 0.0001$).

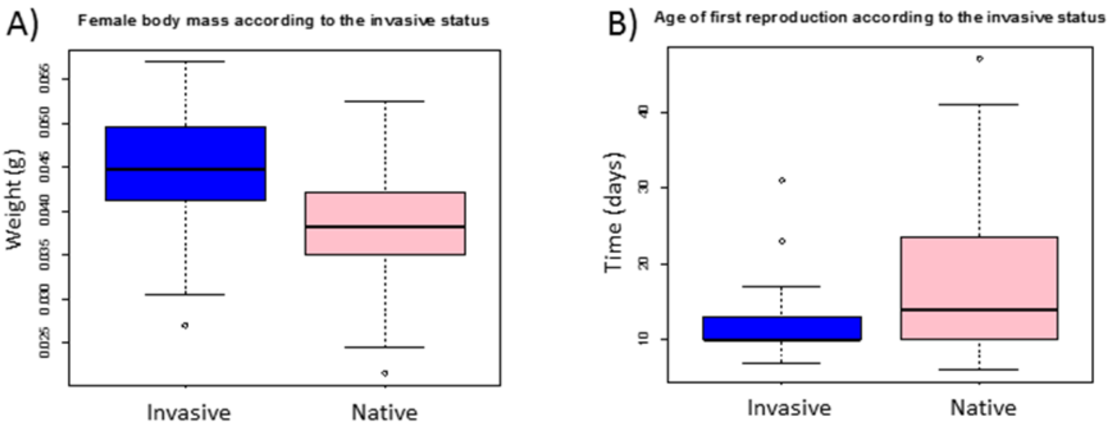

**Supplementary Figure 2:** Founding and selection design for female body mass and age at first reproduction (A) Founding of female body mass lines. (B) Selection of female body mass lines. (C) Founding of age at first reproduction lines. (D) Selection of age at first reproduction lines.

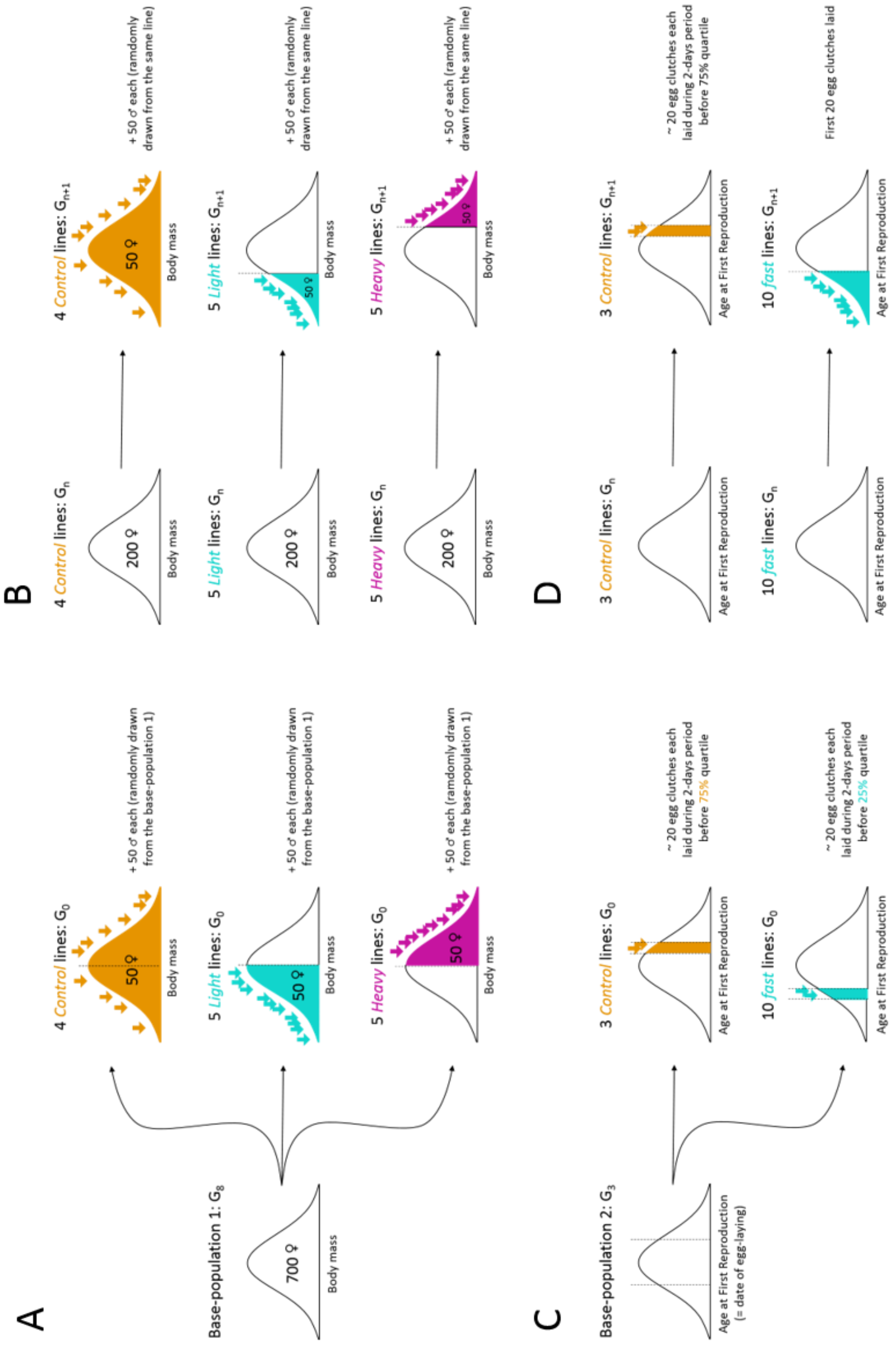

**Supplementary Figure 3: Phenotyping procedure for female body mass (FBM) and age at first reproduction (AFR) lines**

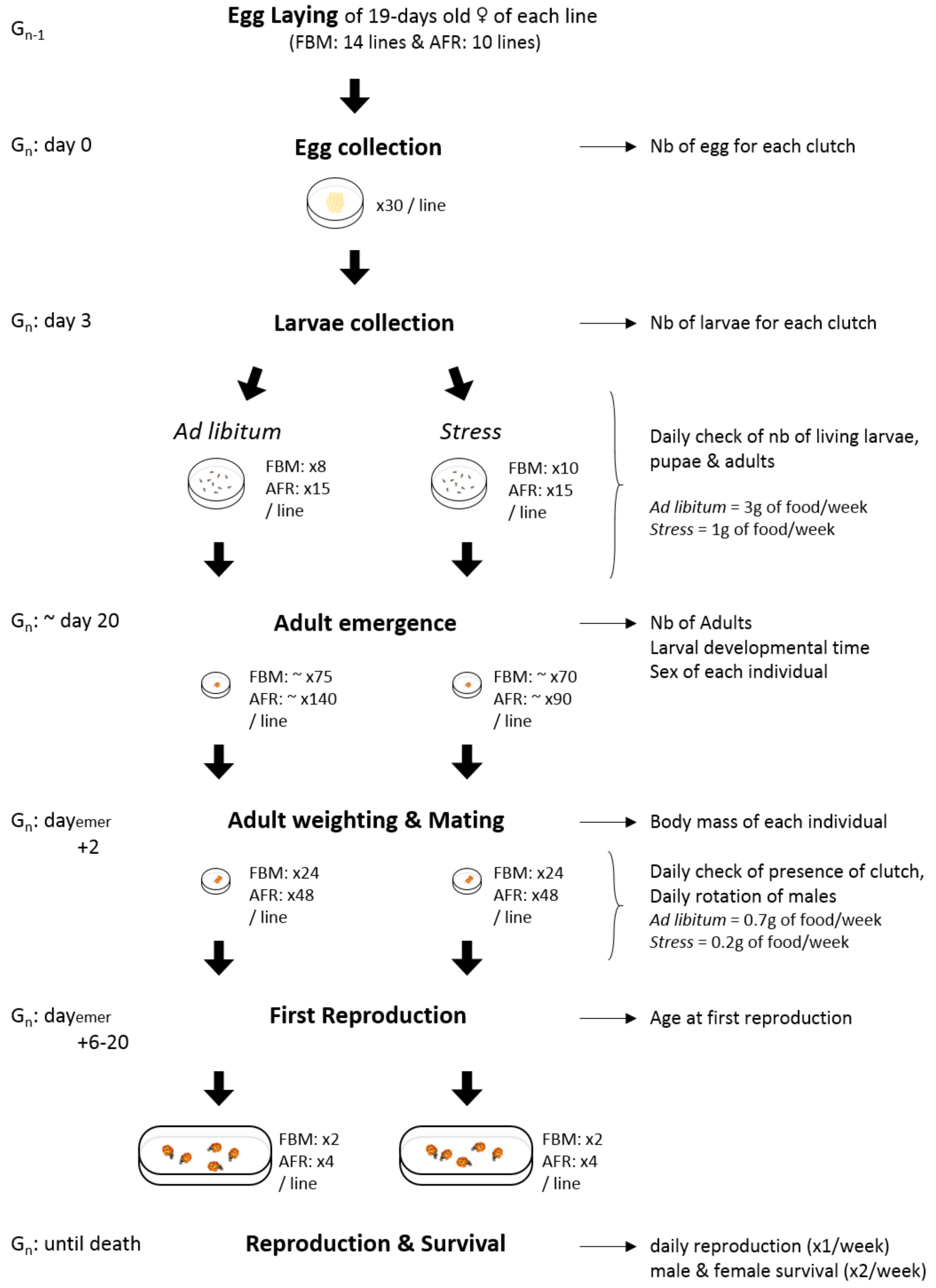

**Supplementary Figure 4: Phenotypic correlation matrices of experimental lines of the female body mass selection scheme**

(A-C) Phenotypic correlation matrices of G9 control, light and heavy lines phenotyped under ad libitum conditions for eight traits. A1R stands for age at first reproduction (D-F) Phenotypic correlation matrices of G9 control, light and heavy lines phenotyped under stressful conditions for eight traits. The phenotypic correlation matrices were strongly dissimilar according to both selection direction and environmental conditions.

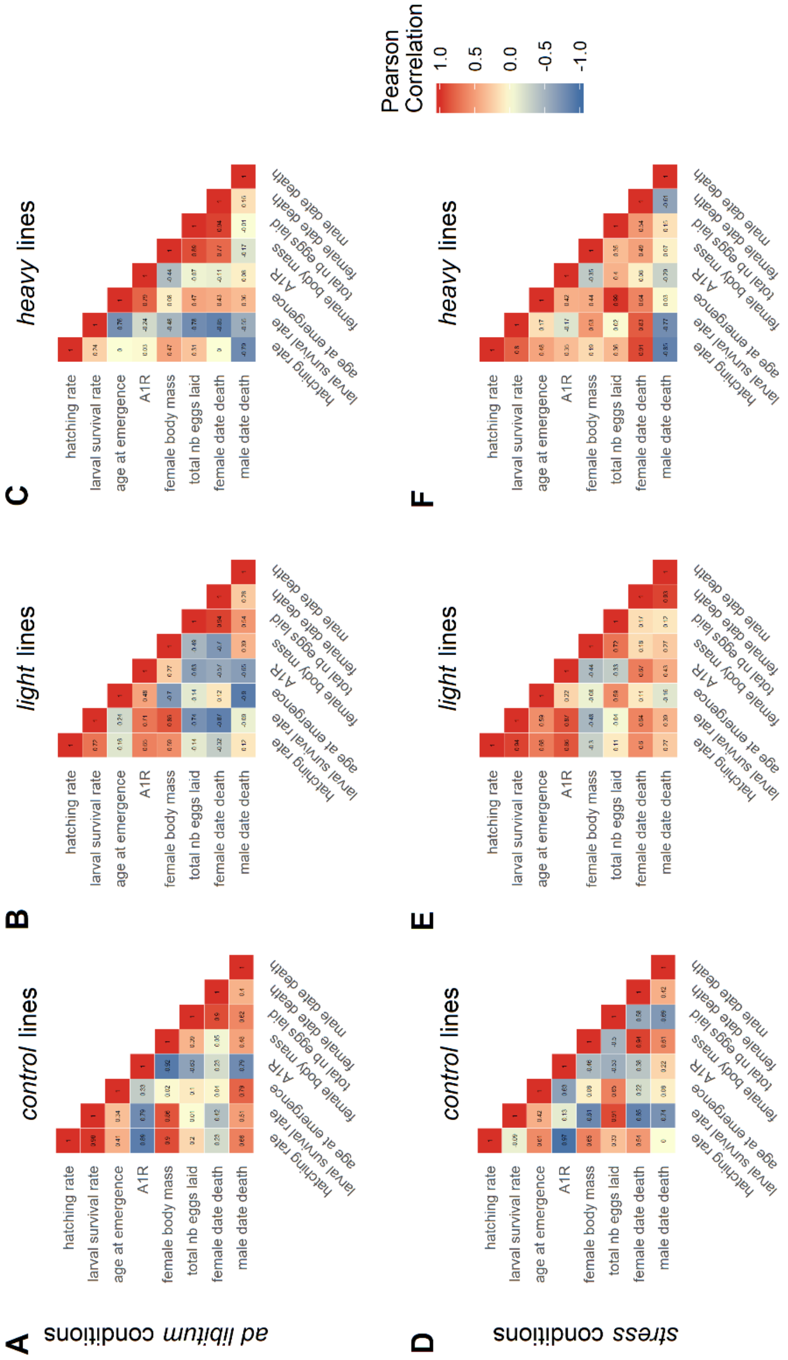

**Supplementary Figure 5: Phenotypic correlation matrices of experimental lines of the age at first reproduction selection scheme**

(A-B) Phenotypic correlation matrices of G10 control and fast lines phenotyped under ad libitum conditions for eight traits. (C-D) Phenotypic correlation matrices of G10 control and fast lines phenotyped under stressful conditions for eight traits. The phenotypic correlation matrices were strongly dissimilar according to both selection direction and environmental conditions. Note also a distinctive pattern of relaxation of phenotypic

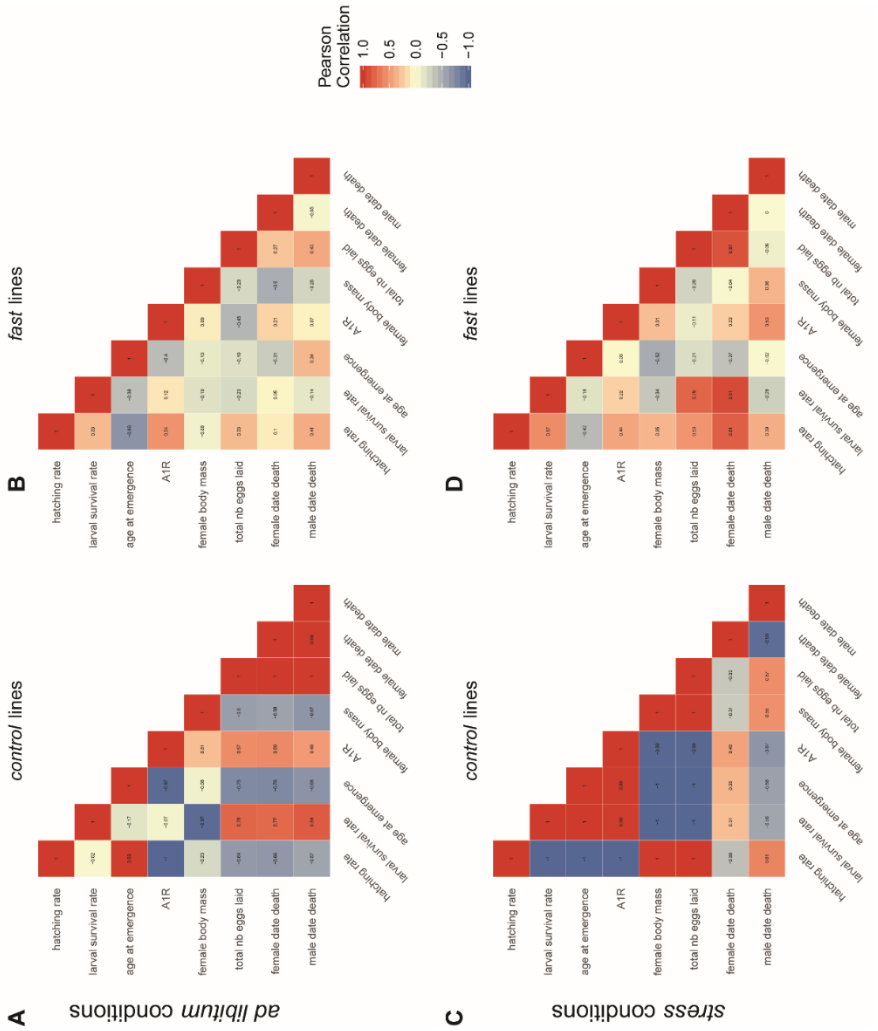

**Supplementary Code & Data:**

Code and data are freely available at the following address: https://data.inra.fr/dataset.xhtml?persistentId=doi:10.15454/V9XCA2

Code file is provided as a .Rmd file (Harmonia Experimental Selection - Code.Rmd)

Description of datasets is given on the repository and in the Rmd code file.
